# Nested parasitism in hypersaline environments: viruses and virus satellites of haloarchaea and their nanosized cellular symbionts

**DOI:** 10.1101/2025.02.15.638363

**Authors:** Yifan Zhou, Ana Gutiérrez-Preciado, Ying Liu, David Moreira, Michail M. Yakimov, Purificación López-García, Mart Krupovic

## Abstract

It is increasingly recognized that hyperparasitism, whereby a parasite exploits a host which itself is a parasite, is a common phenomenon across ecosystems and domains of life. Here, to explore hyperparasitism in Archaea, we focused on ultra-small archaea of the phylum Nanohaloarchaeota, a distinct lineage within the DPANN superphylum, which establish obligate symbiotic interactions with extreme halophiles of the class Halobacteria. We assembled five metagenomes originating from geothermally influenced salt lakes in the Danakil Depression, Ethiopia, and reconstructed the viromes associated with both haloarchaea and nanohaloarchaea. Both archaeal lineages were associated with viruses from four distinct groups, including head-tailed viruses (class *Caudoviricetes*), tailless icosahedral viruses, pleomorphic viruses and spindle-shaped viruses, which represent 12 previously undescribed families. The haloarchaeal viruses (HVs) and nanohaloarchaeal viruses (NHVs) are only distantly related, suggesting that viruses from the four groups co-evolved with their respective hosts for an extended period, likely since the divergence of the two archaeal lineages from their last common ancestor. Consistently, our results show that HVs and NHVs are well-adapted to replicate in their respective hosts and to thrive in hypersaline environments. No evidence of host switching between haloarchaea and nanohaloarchaea was obtained, but multiple horizontal transfers of genes implicated in virion structure and morphogenesis between HVs and NHVs were detected. We also identified several NHVs-encoded auxiliary metabolic genes implicated in nucleotide and amino acid metabolisms, which could enhance the metabolic capabilities of the nanohaloarchaeal hosts that have highly reduced genomes. Finally, in addition to HVs and NHVs, we describe plasmid-derived virus satellites that appear to have originated convergently to parasitize spindle-shaped viruses of both haloarchaea and nanohaloarchaea, uncovering an additional layer of parasitism. Collectively, our findings fill the knowledge gap on the diversity of HVs and NHVs, highlight the complexity of virus-host and virus-virus interactions in hypersaline environments, and open doors for further mining of the virosphere of the globally distributed DPANN archaea.

## Introduction

Outside of the laboratory settings, microbes rarely, if ever, exist in isolation and instead establish complex networks of symbiotic interactions with other cellular organisms and viruses, which range from commensalism and mutualism to parasitism. Remarkably, it has been shown, largely through metagenomics, that a considerable fraction of microbial diversity within both prokaryotic domains is represented by ubiquitous, ultra-small cells with highly reduced genomes, namely, CPR (’Candidate Phyla Radiation’, phylum Patescibacteria)^1^ bacteria and archaea of the DPANN (named after its first representative phyla: Diapherotrites, Parvarchaeota, Aenigmarchaeota, Nanoarchaeota and Nanohaloarchaeota) superphylum^2–4^. Like all cellular organisms, CPR bacteria and DPANN archaea are infected by viruses, but the diversity and impact of these viruses on their hosts remain poorly understood. For instance, it remains unclear whether viruses of CPR bacteria or DPANN archaea underwent genome reduction to adapt to their highly reduced hosts; whether the CPR/DPANN virome has evolved through spillover (host switching) events from the viromes of their respective hosts; whether horizontal gene exchange is common between viruses infecting symbionts and their hosts; whether viruses of symbionts can boost the functional potential of their hosts through auxiliary metabolic genes. Furthermore, it is becoming increasingly recognized that bacterial and archaeal viruses themselves can be targeted by mobile genetic elements, such as virus satellites; however, whether such additional layers of nested parasitism extend to viruses of parasitic hosts, such as CPR bacteria or DPANN archaea, is unknown. To address some of these questions, in this study, we focus on one of the lineages of DPANN archaea, Nanohaloarchaeota, associated with halophilic archaea of the class Halobacteria.

Members of the class Halobacteria and phylum Nanohaloarchaeota represent the dominant archaeal lineages in hypersaline environments^5–8^. Nanohaloarchaea were discovered through metagenomics over a decade ago^9^, but it was not until recently that stable binary cultures including a nanohaloarchaeon and a haloarchaeal host were established under laboratory conditions^10–12^. Culture-based experiments have revealed that nanohaloarchaea have small cells of ∼250-500 nm in diameter and genome sizes of ∼1 Mb^10–12^. The genome reduction is typically accompanied by the loss of metabolic and biosynthetic pathways necessary for energy production and the synthesis of basic biomolecules, such as nucleotides, amino acids and lipids. Accordingly, nanohaloarchaea develop obligate symbiotic relationships with autonomously growing haloarchaea^10–13^. Nanohaloarchaeota often represent a notable fraction of the total archaeal communities in hypersaline habitats^5,9,14^, suggesting that they have evolved efficient strategies for symbiotic lifestyle. One of the mechanisms ensuring their persistence relies on the ability to digest complex polysaccharides, such as glycogen and starch, which may be metabolically inaccessible to their haloarchaeal hosts. However, the simple sugar moieties enzymatically released by nanohaloarchaea can then be readily metabolized by haloarchaea^11^, promoting a mutualistic co-existence of both archaeal groups.

Viruses can affect the composition of microbial communities through host cell lysis and rewire the metabolic capacity of their hosts by supplying diverse auxiliary metabolic genes^15,16^. Hypersaline ecosystems represent some of the most virus-rich environments on the planet^17,18^. Most of the characterized haloarchaeal viruses (HVs) have been discovered through isolation efforts on a relatively small range of host species and are currently classified into 16 families. The majority (n=12) of HVs comprise head-tailed viruses of the class *Caudoviricetes*. However, non-tailed morphotypes are also common in hypersaline environments and include viruses with pleomorphic (*Pleolipoviridae*), tailless icosahedral (*Sphaerolipoviridae* and *Simuloviridae*), and spindle-shaped (*Halspiviridae*) virions^19–21^. Although invaluable for obtaining a comprehensive understanding on the archaeal virosphere, culture-based virus isolation efforts provide a biased view on archaeal virus diversity. Metagenomics allows circumventing the limitations of culture-based virus discovery. For instance, metagenomics provided the first insights into the virome of the square haloarchaeon, *Haloquadratum walsbyi* ^22^, and dramatically expanded the known diversity of haloarchaeal pleolipoviruses^23^. Moreover, metagenomic sequencing allows tracing HV populations across different samples and uncover their dynamics across timescales ranging from days to years^24,25^. While there is a growing appreciation of the diversity and evolution of HVs, little is known about viruses of Nanohaloarchaeota (NHVs), with only of putative NHVs being described through culture-independent approaches^22,24–28^.

Here, we use electron microscopy and metagenomics to explore the diversity of HVs and NHVs in geothermally influenced salt lakes in the Danakil Depression, Ethiopia, some of the most extreme ecosystems known, dominated by microbial communities consisting of haloarchaea and nanohaloarchaea^14,29^. Our results fill the knowledge gap on the diversity and evolution of HVs and NHVs, and illuminate the complexity of nested virus-host and virus-virus interactions.

## RESULTS

### Virus-like particles in samples from Danakil salt lakes

Microbial communities in hypersaline environments are dominated by archaea and are under constant attack by diverse viruses^30^. To explore the viromes of consortia consisting of haloarchaea (class Halobacteria) and symbiotic nanohaloarchaea (phylum Nanohaloarchaeota), we focused on salt lake samples from the Danakil Depression, Ethiopia, a region with multiple hypersaline systems of diverse hydrochemistry (Fig. 1A)^14,29^. Haloarchaea and nanohaloarchaea were dominant in the samples from Lake Assale or Karum (samples Ass and 9Ass collected during different years), cave reservoir at the Dallol proto-volcano salt canyons (9Gt) and two of the Western-Canyon Lakes (WCL2 and WCL3) (Fig. 1B)^29^. Transmission electron microscopy analysis of the enrichment cultures established from the 9Ass, 9Gt and WCL3 samples revealed the presence of diverse virus-like particles (VLPs; Fig. 1C). In the 9Ass and 9Gt enrichments only spherical VLPs were observed. By contrast, the VLPs in the WCL3 enrichment were more diverse (Fig. 1C), including head-tailed VLPs typical of the class *Caudoviricetes*, with non-contracted and contracted tails, as well as several types of spherical VLPs. The different types of spherical VLPs could be distinguished based on the diameter and different staining properties (Fig. 1C). Both head-tailed and spherical VLPs are commonly observed in hypersaline environments^19,31^. These results confirmed the presence of diverse viruses in the Danakil salt lake samples.

**Figure 1.**
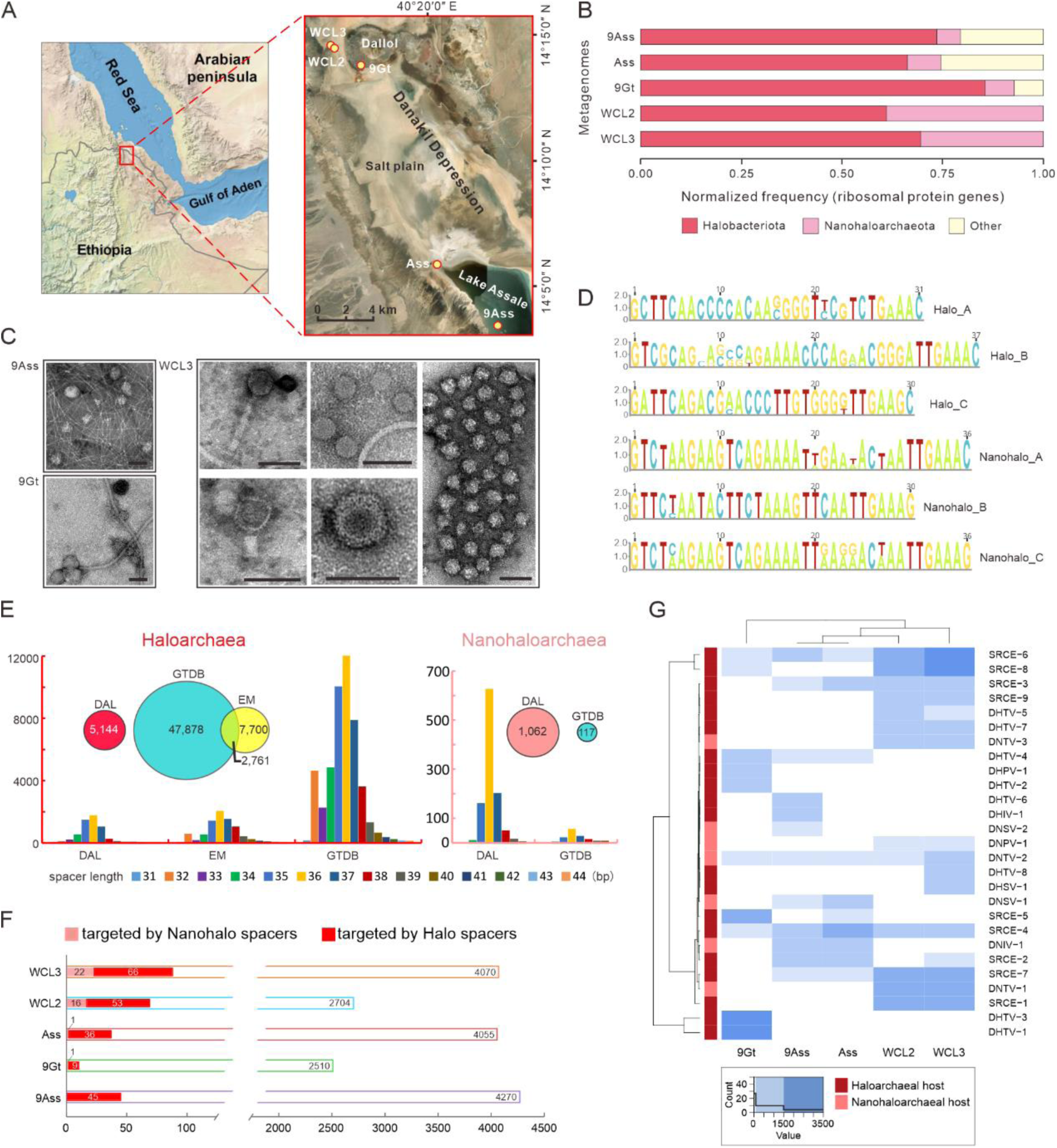
Microbial and viral diversity in the salt lakes of the north Danakil Depression, Ethiopia. a. Locations of the five sampling sites in the Danakil Depression. b. Microbial community composition in the environmental samples collected from the five salt lakes. The composition is inferred from the previously described normalized frequency of selected ribosomal proteins ^29^. c. Electron micrographs of virus-like particles observed in the enrichment cultures established from environmental samples collected from the 9Ass, 9Gt and WCL3 salt lakes. The virus-like particles were stained with 2% (w/v) uranyl acetate. Scale bars, 100 nm. d. The consensus sequences of the six CRISPR variants specific to Halobacteriota and Nanohaloarchaeota. e. The total numbers and length distributions of spacers extracted from the Danakil Depression (DAL) metagenomes, Earth Microbiome (EM) Project and Genome Taxonomy Database (GTDB). f. The results of CRISPR spacer targeting. The open bars represent the total number of viral sequences identified in each metagenome, whereas the filled bars represent the number of viral sequences targeted by Halo (red) or Nanohalo (pink) spacers. g. Heatmap showing the distribution and abundance of haloarchaeal and nanohaloarchaeal viruses and plasmid-like elements in Danakil salt lakes.

### Overall composition of the Danakil viromes

To get an unbiased view on the diversity of HVs and NHVs in the Danakil salt lake samples, five metagenomes were analyzed for the presence of viral contigs using geNomad ^32^ and VirSorter2 ^33^, which collectively yielded 2,085 viral contigs (≥5 kb). The vConTACT gene-sharing network analysis showed that the majority (n=1641) of Danakil viral sequences (including 950 singletons) were not connected to any of the reference prokaryotic viruses and thus represent novel viruses (Fig. S1). The contigs that were related to known viruses were mainly connected to three archaeal virus clusters: i) head-tailed HVs (class *Caudoviricetes*, n=382); ii) tailless icosahedral HVs (families *Simuloviridae* and *Sphaerolipoviridae*, n=26); iii) pleomorphic HVs (family *Pleolipoviridae*, n=9) (Fig. S1). Only 1% (21/2,085) of viral sequences were related to bacteriophages, consistent with the fact that archaea are predominant members of the prokaryotic communities in the Danakil salt lakes (Fig. 1B). However, we note that the abundance of bacterial viruses may be underestimated due to scarcity of halophilic bacteriophage reference genomes. Regardless, these results highlight the novelty of the Danakil viromes and suggest that viruses infecting halophilic archaea are the major players in these hypersaline environments.

### Haloarchaea and nanohaloarchaea are associated with distinct viromes

Both Halobacteria and Nanohaloarchaeota species carry CRISPR arrays^13,34^. Thus, the CRISPR spacers within these arrays can be used to trace the historical encounters between the corresponding archaeal species and mobile genetic elements, including viruses. To link the identified virus contigs to their potential hosts, we assembled a database of CRISPR sequences specific to haloarchaeal or nanohaloarchaeal species (see Methods). A total of 64 haloarchaea-specific CRISPR sequences and 11 nanohaloarchaea-specific CRISPR sequences were identified in the Danakil metagenomes, which together with related CRISPR arrays from the Genome Taxonomy Database (GTDB) and Earth’s Microbiomes (EM) dataset^35^, formed 6 distinct clusters (Fig. 1D and Fig. S2).

The haloarchaea-specific CRISPR arrays from the Danakil metagenomes, GTDB and EM contained 5,144, 47,878 and 7,700 spacers, respectively (Fig. 1E). In the case of nanohaloarchaea, 1,062 and 117 spacers were extracted from Danakil metagenomes and GTDB, respectively, whereas EM did not contain identifiable nanohaloarchaeal spacers. Spacers extracted from different databases displayed similar length distributions, centering at ∼36 bp (Fig. 1E). The Danakil haloarchaeal and nanohaloarchaeal spacers showed no overlap with those from the two other databases, further pointing to the novelty of the Danakil viral assemblages.

The collected CRISPR spacers were used as queries to search against the Danakil viral contigs (≥1 kb). A total of 209 and 44 viral contigs were targeted (≥30 bp exact match) by haloarchaeal and nanohaloarchaeal spacers, respectively (Fig. 1F). The spacer-targeted sequences represented only a very small fraction of the whole virome (<2%). Notably, no sequences were found to be targeted by both nanohaloarchaeal and haloarchaeal spacers. Although the number of CRISPR-targeted haloarchaeal and nanohaloarchaeal virus genomes remains limited, the obtained results suggest that there is no overlap between the corresponding viromes, despite the intimate interaction between the two archaeal lineages.

### Diversity and distribution of haloarchaeal and nanohaloarchaeal viruses

Following contig extension (see Methods), we obtained 11 and 6 complete genomes of Danakil HVs and NHVs, respectively (Table S1). Based on the presence of signature genes involved in virion morphogenesis, these viruses could be broadly assigned to four realm-level groups (highest rank in virus taxonomy), namely, (i) head-tailed viruses of the realm *Duplodnaviria* (class *Caudoviricetes*; 8 HVs and 3 NHVs), (ii) tailless icosahedral viruses of the realm *Singelaviria* (formerly a kingdom within realm *Varidnaviria*; 1 HVs and 1 NHVs), (iii) pleomorphic viruses of the realm *Monodnaviria* (family *Pleolipoviridae*; 1 HVs and 1 NHVs), and (iv) spindle-shaped viruses which are currently not assigned to any realm and likely to constitute a realm of their own in the future (1 HVs and 1 NHVs as well as 1 near-complete NHV genome). Hereinafter, we refer to Danakil haloarchaeal and nanohaloarchaeal tailed viruses as DHTVs and DNTVs; to tailless icosahedral viruses as DHIVs and DNIVs; to pleomorphic viruses as DHPVs and DNPVs; and spindle-shaped viruses as DHSVs and DNSVs, respectively. In addition to viral genomes, we identified 9 spacer-targeted small plasmid-like circular elements associated with both haloarchaea and nanohaloarchaea (see below).

Analysis of the distribution of HVs and NHVs across the five Danakil salt lake metagenomes revealed that the viromes varied across the studied ecosystems, with the patterns for the cave reservoir (9Gt), the Lake Assale samples (Ass, 9Ass) and the Western Canyon Lakes (WCL2, WCL3) being clearly distinct (Fig. 1G). Overall, all lakes contained both HVs and NHVs, but certain viruses were exclusive to particular lakes. For instance, DHTV1-3 and DHPV-1 were restricted to 9Gt, whereas DNSV−1, DNSV-2, DNIV-1, DHIV-1, and DHTV-6 were only found in Lake Assale. However, some of the viruses, in particular, DNTVs and DHTVs, and plasmid-like elements were found across several lakes, with DNTV-2 being detectable across all sites (Fig. 1G). This differential virome composition is likely shaped by both abiotic (physicochemical parameters, including differences in chaotropicity) and biotic (host) environmental selection.

### Deep divergence of haloarchaeal and nanohaloarchaeal viromes

To further explore the diversity of NHVs, we searched the IMG/VR and NCBI databases for the presence of virus genomes related to those identified in the Danakil metagenomes (see Methods). These searches yielded additional complete or near-complete genomes related to DNTVs (n=7), DNIV-1 (n=3), DNPV-1 (n=1) and DNSVs (n=1; Table S2, Fig. S3, Fig. S4). Host assignments for all complete genomes were done using a combination of CRISPR targeting and blastp searches for proteins matching the reference cellular or viral proteomes (Table S1 and S2; see Methods).

Comparison of the inferred proteomes of HVs and NHVs showed that the corresponding virus groups are only distantly related to each other (Fig. 2), arguing against recent horizontal virus transfer between haloarchaeal and nanohaloarchaeal hosts. In particular, proteome-wide comparison of DHTVs and DNTVs in the context of the reference archaeal viruses representing diverse *Caudoviricetes* families showed that DHTVs and DNTVs formed distinct families (Fig. 2A). Based on the established demarcation criteria, i.e., archaeal viruses of the same *Caudoviricetes* family typically share about 20-50% of orthologous proteins, whereas viruses from different families share less than 10% of proteins^36^, the 8 DHTVs could be classified into 7 different families, including 4 new families, whereas the DNTVs and their relatives formed 4 new families (Fig. S5, Fig. S6 and Table S1). These assignments were consistent with the proteomic tree analysis, where the branch length demarcation for different archaeal *Caudoviricetes* families is ∼0.05. Notably, members of the same family exhibited similar genome lengths and GC% content (Fig. 2), further supporting the taxonomic assignments.

**Figure 2.**
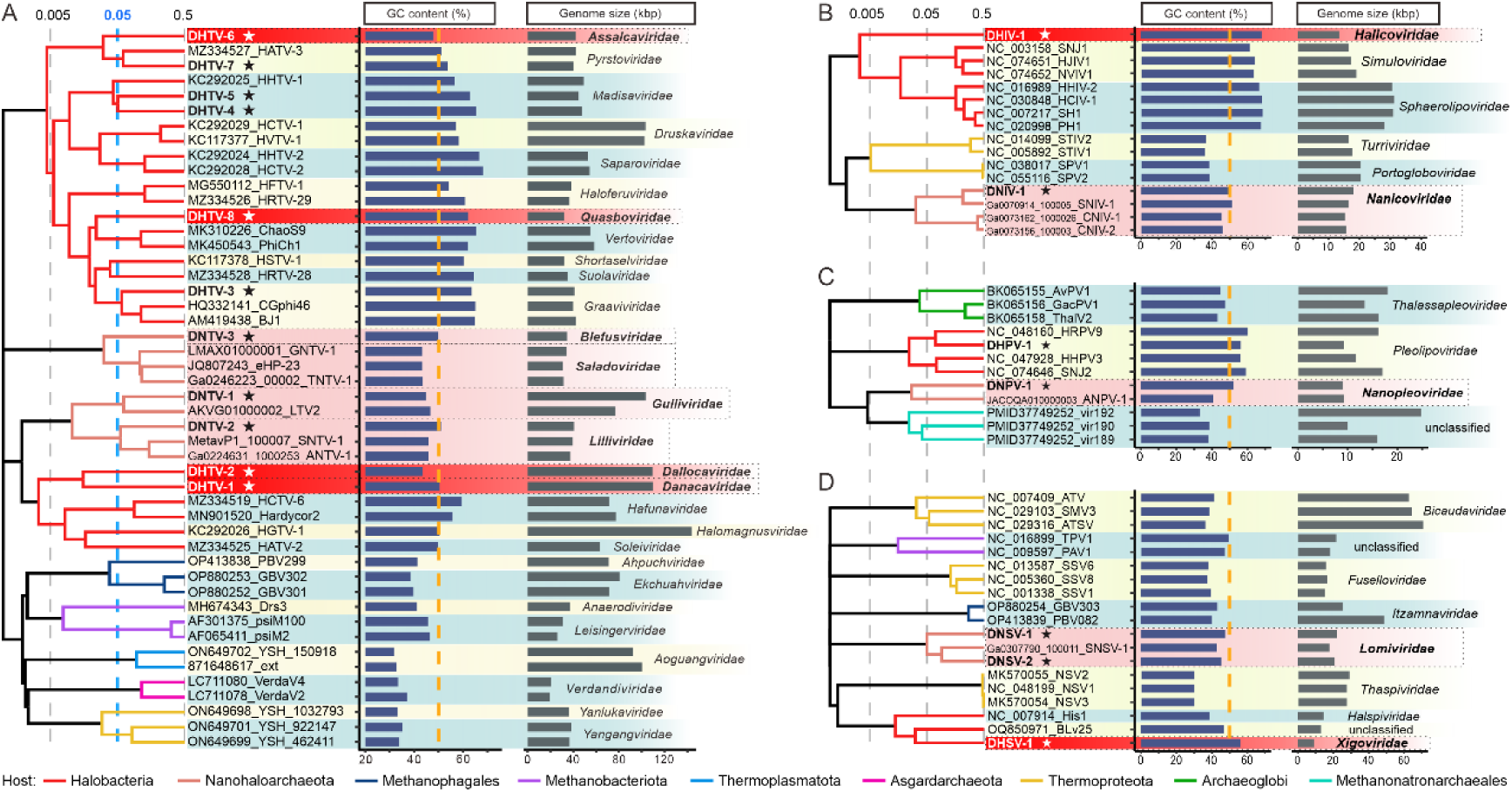
Genome-wide proteomic trees of the four virus groups. The bars next to each genome indicate the GC% content and genome length, respectively. a. Head-tailed viruses (*Caudoviricetes*). b. Tailless icosahedral viruses. c. Pleomorphic viruses. d. Spindle-shaped viruses. Viruses from the Danakil Depression are indicated with stars. Proposed new families are highlighted in red (HVs) or pink (NHVs). The proteomic trees are based on all-versus-all proteomic similarity matrix and are mid-point rooted. Branch lengths are log-scaled. Branches are colored based on the viral host groups, with the key provided at the bottom of the figure.

The proteomic trees showed that other HVs and NHVs also formed distinct family-level groups, with only pleomorphic DHPV-1 (9.2 kb) falling within the existing haloarchaeal virus family *Pleolipoviridae* (Fig. 2C and Fig. S7A). By contrast, icosahedral DHIV-1 (13.6 kb) and DNIV-1 (18 kb) as well as pleomorphic DNPV-1 (9.1 kb), together with the relatives retrieved from the public databases, each formed potential new families (Fig. 2B, Fig. 2C, Fig. S7, Fig. S8). DHIV-1 and DNIV-1 are only distantly related to tailless icosahedral HVs of the family *Simuloviridae*, with the similarity between the corresponding major capsid proteins (MCPs) being detectable only at the level of sensitive profile-profile comparisons.

We detected a provirus closely related to DNPV-1 integrated within the genome of a nanohaloarchaeon (JACOQA010000003) from the Atlit salt pools in Israel ^7^, which we refer to as ANPV-1 (Fig. 2C and Fig. S7B). Although haloarchaeal and nanohaloarchaeal pleomorphic viruses encode a canonical morphogenetic module typical of this group and including homologs of the HRPV1 matrix protein VP3, membrane fusion protein VP4, AAA+ ATPase as well as ORF6 and ORF7, the similarity between the DHPV-1 and DNPV-1 homologs is not detectable by blastp (Fig. S7 and Fig. 3). Accordingly, DNPVs cluster outside of the haloarchaeal *Pleolipoviridae*, forming a sister group to the unclassified pleomorphic viruses of Methanonatronarchaeales, an order of halophilic methanogens ^37,38^. Notably, whereas DHPV-1 encodes a rolling circle replication initiation endonuclease (Fig. S7 and Fig. 3), typical of viruses in the genus *Alphapleolipovirus*, its morphogenetic module is more closely similar to those of betapleolipoviruses, suggesting that DHPV-1 might represent a new genus within the family. By contrast, DNPV-1 and ANPV-1 do not encode detectable replication proteins.

**Figure 3.**
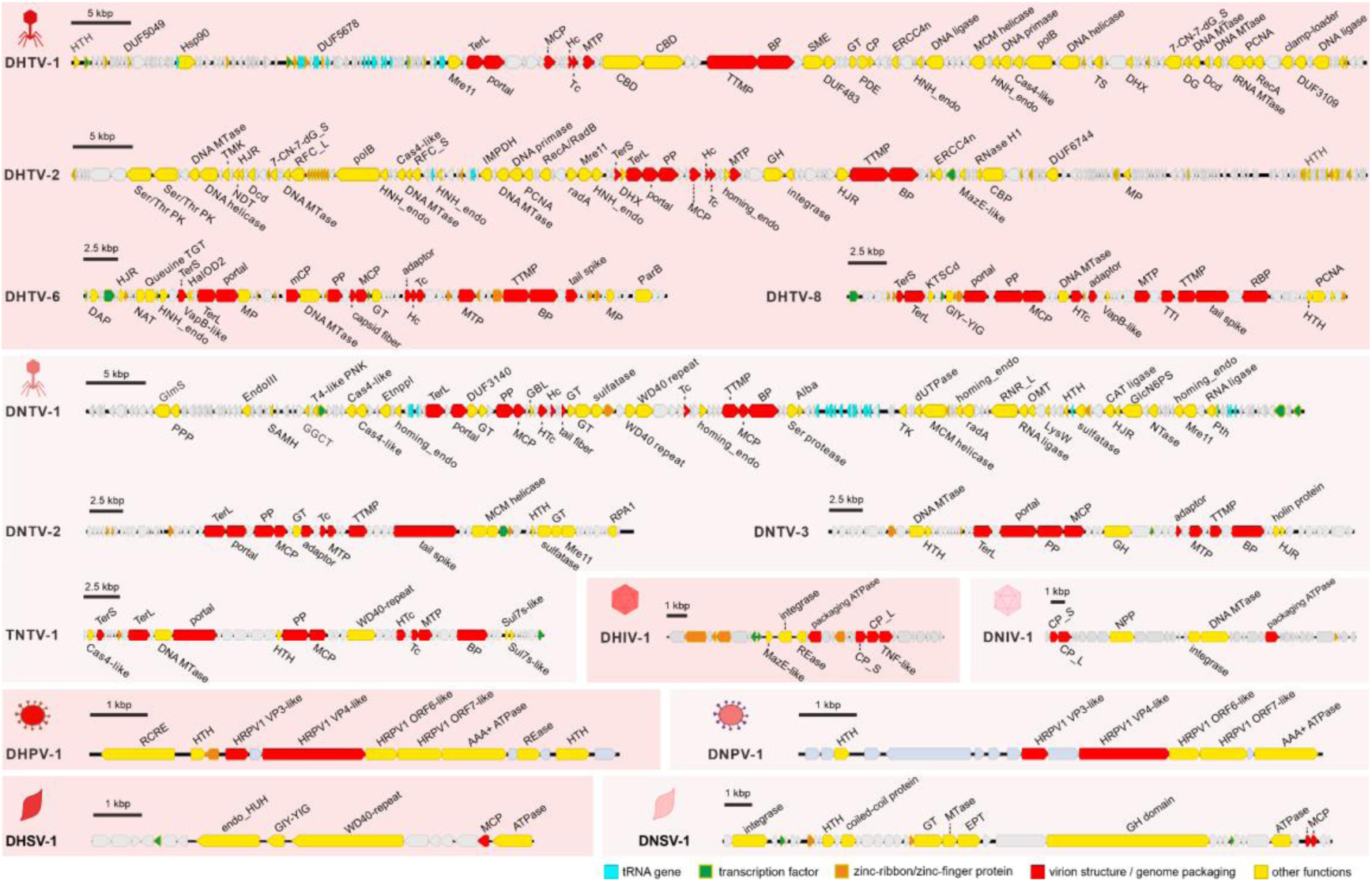
Genome maps of HVs and NHVs from Danakil Depression. For head-tailed viruses, only representatives of the proposed new families are shown. Each family is represented by a single genome. Abbreviations: HTH, helix-turn-helix; DUF, domain of unknown function; Hsp90, ATP-dependent chaperone Hsp90; Mre11, nuclease Mre11; TerS and TerL, small and large subunits of the terminase, respectively; MCP, major capsid protein; Hc, head-closure protein; Tc, tail-completion protein; MTP, major tail protein; CBD, carbohydrate binding domain; TTMP, tail tape measure protein; BP, baseplate protein; SME, sulfatase-maturating enzyme; GT; glycosyltransferase; PDE, phosphodiesterase; CP, cysteine protease; ERCC4n, ERCC4-type nuclease; HNH_endo, HNH family endonuclease; polB, family B DNA polymerase; DHX, DEAD/DEAH-box helicase; 7-CN-7-dG_S, 7-cyano-7-deazaguanine synthase; DNA MTase, DNA methyltransferase; DG, DNA glycosylase; Dcd, dCTP deaminase; tRNA MTase, tRNA methyltransferase; PCNA, DNA polymerase sliding clamp; RecA, RecA ATPase; Ser/Thr PK, serine/threonine protein kinase; TS, thymidylate synthase; TMK, thymidylate kinase; TK, thymidine kinase; HJR, Holliday junction resolvase; NDT, nucleoside 2-deoxyribosyltransferase; RCF_L, replication factor C large subunit; RCF_S, replication factor C small subunit; IMPDH, inosine-5’-monophosphate dehydrogenase; RecA/RadB-like, RecA/RadB-like recombination protein; RadA, DNA repair and recombination protein RadA; PP, prohead protease; homing_endo, homing endonuclease; GH, glycoside hydrolase; RNase H1, ribonuclease HI; MazE-like, MazE-like antitoxin; CBP, cobalamin biosynthesis protein; MP, metalloprotease; DAP, DNA annealing protein; NAT, N-acetyltransferase; Queuine TGT, queuine tRNA-guanine transglycosylase; VapB-like, VapB-like antitoxin; HalOD2, haloarchaeal output domain 2; mCP, minor capsid protein; ParB, ParB family DNA-binding protein; KTSCd, lysine (K) tRNA synthetase C-terminal domain; GIY-YIG, GIY-YIG family nuclease; HTc, head-tail connector protein; TTI, tail tube initiator; RBP, receptor binding protein; GlmS, glucosamine 6-phosphate synthase; PPP, phosphotyrosine protein phosphatase; EndoIII, endonuclease III; SAMH, S-adenosyl-L-methionine hydrolase; GGCT, gamma-glutamyl cyclotransferase; T4-like PNK, T4-like polynucleotide kinase; Etnppl, ethanolamine phosphate transferase; GBL, galactose-binding lectin; WD40 repeat, WD40 repeat-containing protein; Alba, DNA/RNA-binding protein Alba; RNR_L, ribonucleotide reductase large subunit; OMT, O-methyltransferase; LysW, amino group carrier protein LysW; GlcN6PS, glucosamine 6-phosphate synthetase; CAT ligase, carboxylate-amine/thiol ligase; Pth, peptidyl-tRNA hydrolase; PPA1, archaeal replication protein A1; Sul7s-like, Sul7s-like DNA binding protein; CP_S, capsid protein (small); CP_L, capsid protein (large); REase, restriction endonuclease; TNF-like, TNF-like jelly-roll domain protein; NPP, nucleotide pyrophosphatase/phosphodiesterase; RCRE, rolling circle replication endonuclease; HRPV1 VP3-like, HRPV1 VP3-like matrix protein; HRPV1 VP4-like, HRPV1 VP4-like membrane fusion protein; endo_HUH, endonuclease of the HUH superfamily; EPT, ethanolamine phosphate transferase.

The putative spindle-shaped viruses of haloarchaea and nanohaloarchaea, DHSV-1 (9.0 kb) as well as DNSV-1 (22.1 kb) and DNSV-2 (20.8 kb), respectively, that encode the signature hydrophobic α-helical hairpin MCP^39,40^, likely also form two distinct families (Fig. 2D and Fig. S3). Notably, the morphogenetic module of DHSV-1, including the MCP and AAA+ ATPase, is most closely similar to those of *Halorubrum* spindle-shaped virus BLv25 and *Haloarcula hispanic*a virus His1 (family *Halspiviridae*) (Fig. S3B and Fig. 3). However, unlike BLv25 and His1, DHSV-1 does not encode a protein-primed family B DNA polymerase and instead codes for an endonuclease of the HUH superfamily that is responsible for rolling circle replication initiation (Fig. S3B and Fig. 3). This finding underscores the diversity of genome replication mechanisms among spindle-shaped HVs. The morphogenetic module of the three DNSVs consists of AAA+ ATPase and two paralogous MCPs (Fig. S3A and Fig. 3), a characteristic previously observed in spindle-shaped viruses of Asgard archaea (family *Wyrdviridae*)^41^. Notably, DNSVs share an integrase of the tyrosine recombinase superfamily, suggesting that they are capable of lysogenizing their nanohaloarchaeal hosts, which provides for vertical replication of the viral genome. DNSV-1 and DNSV-2 both encode a large gene (>1900 codons) containing glycoside hydrolase domain (Fig. S3A) which is likely to be the receptor-binding adhesin as found in virions of hyperthermophilic fuselloviruses ^39^. However, as is typical with archaeal virus genomes ^42^, most of the DNSV genes cannot be functionally annotated, and genes for genome replication cannot be predicted.

Collectively, these results suggest that HVs and NHVs can be classified into 16 families, 12 of which have not been described previously, emphasizing sparse sampling of this part of the archaeal virome. Notably, none of the HVs and NHVs displayed sufficient sequence similarity to be considered as members of the same family, suggesting deep divergence of the corresponding viromes.

### Amino acid composition of HVs and NHVs reflects adaptation to high salt environments

To thrive under close-to-saturating salt concentrations, nanohaloarchaea and haloarchaea both have adopted the “salt-in” strategy, whereby the intracellular salt concentration is similar to that in the surrounding environment ^26,29,43^. One of the most prominent molecular adaptations permitting the “salt-in” strategy is the prevalence of negatively charged amino acids in the haloarchaeal proteomes, which ensures the proper folding and stability of the corresponding proteins^43^. Notably, the most acidic proteomes (median pI≤4.4) ever observed are encoded by archaea from the salt lakes of Danakil depression, specifically, those from the Western-Canyon Lakes^29^. However, whether HVs and NHVs display similar adaptations to intracellular and extracellular high salt environments has not been explicitly addressed. Therefore, we examined the amino acid usage (AAU) patterns and proteome-wide isoelectric point (pI) distributions of HVs and NHVs, and compared them with those of the respective predicted hosts (Fig. S9A).

The proteomes of haloarchaea and nanohaloarchaea have independently adapted to high salt environments and display differences in the prevalence of certain amino acids^6^. Most prominently, haloarchaeal and nanohaloarchaeal proteomes tend to be strongly enriched in alanine (A) and glutamate (E) residues, respectively. The same AAU patterns were found in the viruses predicted to infect the two groups of halophilic archaea. More generally, the enrichment of the HV and NHV proteomes in negatively charged amino acid residues (D and E) yields similar pI distributions for viruses and their hosts, which are distinct from those of non-halophilic archaea (Fig. S9B).

Adaptation of HVs and NHVs to their hosts is also evident at the level of nucleotide composition. Analysis of the GC% content showed that nanohaloarchaea (42.9%; n=38) have considerably lower GC% content compared to that of haloarchaea (64.1%; n=749) and the same was true for head-tailed viruses predicted to infect the two host groups (58.2% [n=62] versus 46.1% [n=9] for DHTVs and DNTVs, respectively; Fig. S9C).

These patterns suggest that HVs and NHVs have adapted their AAU and GC contents to mirror those of their hosts, presumably to optimize the compatibility with the intracellular pools of amino acids and nucleotides. The distinct GC and AAU patterns further suggest that haloarchaeal and nanohaloarchaeal viromes have diverged early on and coevolved with their hosts for an extended period. Conversely, the disparity between HVs and NHVs argues against the possibility that either group of viruses can infect both hosts, consistent with the lack of viral genomes that would be targeted by CRISPR spacers from both haloarchaeal and nanohaloarchaea.

### Virus auxiliary metabolic genes can enhance the hosts’ metabolic potential

Viruses, especially those with larger genomes, often carry auxiliary metabolic genes (AMG), which can supplement and enhance the functional potential of the host cells. Although nanohaloarchaea have dramatically reduced genomes, their viruses do not appear to follow the same trend. Indeed, HVs and NHVs displayed similar genome sizes, with the largest viral genomes in both host groups exceeding 100 kb (DHTV-1 with 109.7 kb versus DNTV-1 with 103.4 kb).

Most of the genes (among those which could be functionally annotated) in the HVs and NHVs with smaller genomes (<50 kb) encoded the core functions involved in virion morphogenesis and genome replication, whereas viruses with large genomes encode more diverse functions. For instance, DNTV-1 and DNTV-2 encode nearly complete replisomes and a number of AMGs, particularly those involved in nucleotide metabolism (see Supplementary text). By contrast, the similarly sized genome of DNTV-1 lacks the expansive set of genome replication proteins and only encodes the small subunit of the sliding clamp loader and MCM helicase (Fig. 3 and Table S3). The latter protein is commonly found as a sole replication protein in archaeal viruses with smaller genomes^44^. Consistently, in the proteomic tree (Fig. 2A), DNTV-1 clusters with smaller DNTVs, suggesting that this virus underwent genome expansion independently from tailed viruses of halophilic and marine archaea.

Although nanohaloarchaea largely depend on haloarchaea for the supply of amino acids and nucleotides, similar to DHTV-1 and DHTV-2, DNTV-1 encodes several proteins implicated in nucleotide metabolism, including deoxyuridine triphosphatase (ORF88) responsible for the hydrolysis of dUTP to pyrophosphate and dUMP, which can then be used for the synthesis of dTMP; thymidine kinase (ORF91), which phosphorylates (d)T to (d)TMP; and the large subunit of the ribonucleoside-diphosphate reductase (ORF77; RNR), which uses thioredoxin (ORF93) as a cofactor to convert ribonucleotides into deoxyribonucleotides (Fig. 3 and Table S3). A relative of DNTV-1, LTV2, a virus found in hypersaline Lake Tyrrell, Australia but not previously linked to nanohaloarchaeal hosts ^25^, also encodes an RNR (Fig. S4A and Table S3). Although DNTV-1 and LTV2 are assigned to the same family (Fig. 2A, Fig. S5), their RNRs are not orthologous but have been independently acquired horizontally from nanohaloarchaeal hosts (Fig. 5A, see Supplementary text). It is tempting to speculate that NHVs played an active role in the spread of the *RNR* genes in the archaeal populations.

In addition, DNTV-1 carries several genes implicated in translation and amino acid metabolism. In particular, it encodes 29 tRNAs for 19 different amino acids (Table S4), including 5 tRNAs for the negatively charged Asp and Glu residues that are enriched in the nanohaloarchaeal and NHV proteomes (Fig. S9A). Other translation related genes include archaeal-type peptidyl-tRNA hydrolase (ORF50; Fig. 3, Fig. 4B), an enzyme that cuts unfinished peptides off tRNAs that have been prematurely released from a stalled ribosome^45^ and two copies of phage T4-like RNA ligase 2 (ORF51 and ORF76, Fig. 3 and Table S3), which is implicated in tRNA repair ^46^. DNTV-1 also encodes a GGCT/AIG2 family gamma-glutamyl cyclotransferase (ORF158, Fig. 4B), an enzyme catabolizing L-γ-glutamyl-L-α-amino acid dipeptides formed during the γ-glutamyl cycle to 5-oxo-L-proline and a free amino acid ^47^.

**Figure 4.**
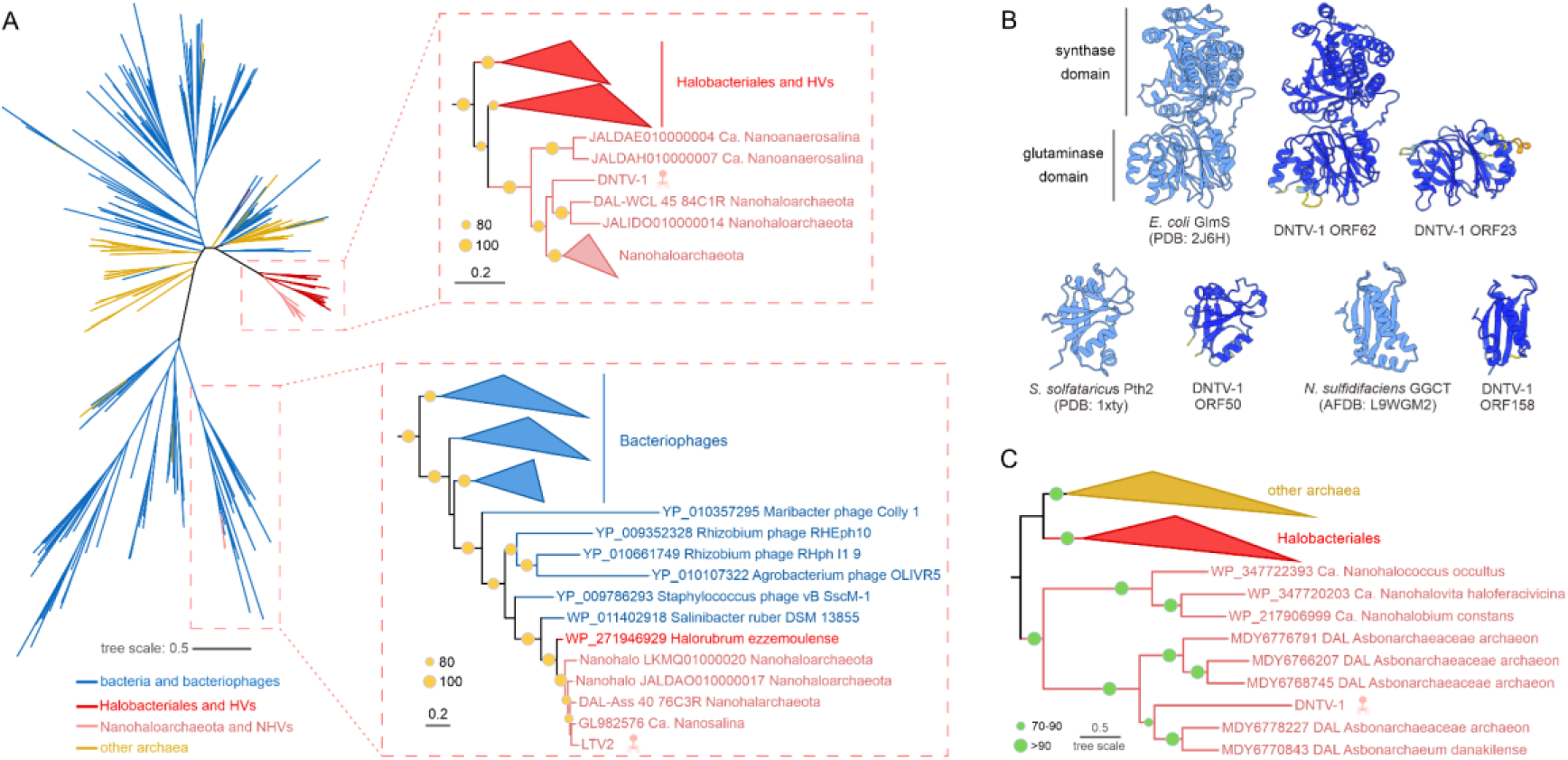
Phylogenetic analysis and structural modeling of AMGs encoded by DNTV-1. a. Unrooted maximum likelihood phylogeny of ribonucleotide reductase large subunit. Zoom-ins on the branches including DNTV-1 and LTV2 are shown on the right. Branches with bootstrap support values higher than 80% are indicated with yellow circles. b. Predicted structural models of DNTV-1 encoded glucosamine 6-phosphate synthase (GlmS; ORF62), standalone GlmS glutaminase domain (ORF23), archaeal-type peptidyl-tRNA hydrolase (Pth2; ORF50), GGCT/AIG2 family gamma-glutamyl cyclotransferase (ORF158). c. Mid-point rooted maximum likelihood phylogeny of glucosamine 6-phosphate synthase (GlmS). Branches with bootstrap support values higher than 70% are indicated with green circles.

Finally, DNTV-1 ORF62 encodes a glucosamine 6-phosphate synthase (GlmS), an enzyme which operates at a crossroad between amino acid and carbohydrate metabolism and catalyzes conversion of Gln and fructose-6-phosphate to Glu and glucosamine-6-phosphate ^48^. The protein contains two domains (Fig. 4B), an N-terminal glutaminase domain, which catalyzes the hydrolysis of Gln to Glu and ammonia, and a C-terminal synthase domain, catalyzing amination of fructose-6-phosphate. In addition, DNTV-1 ORF23 encodes a stand-alone GlmS glutaminase domain (Fig. 5B)^49^, suggesting that it participates in Gln deamination but not carbohydrate amination. This activity could increase the pool of intracellular Glu, one of the most prevalent amino acids in the proteomes of NHVs and nanohaloarchaea (Fig. S9A). In a phylogeny, DNTV-1 GlmS branches within a clade containing nanohaloarchaeal homologs and is closest to proteins encoded by members of the family Asbonarchaeaceae (Fig. 4C), a basal lineage of Nanohaloarchaeota recently described in Danakil salt lakes ^6^. In addition, four other DNTV-1 proteins yielded best blastp hits to Asbonarchaeaceae (Table S8). Notably, DNTV-2 was binned into the Asbonarchaeaceae MAG. These results suggest that DNTV-1 and DNTV-2, and potentially other NHVs infect autochthonous members of the family Asbonarchaeaceae.

**Figure 5.**
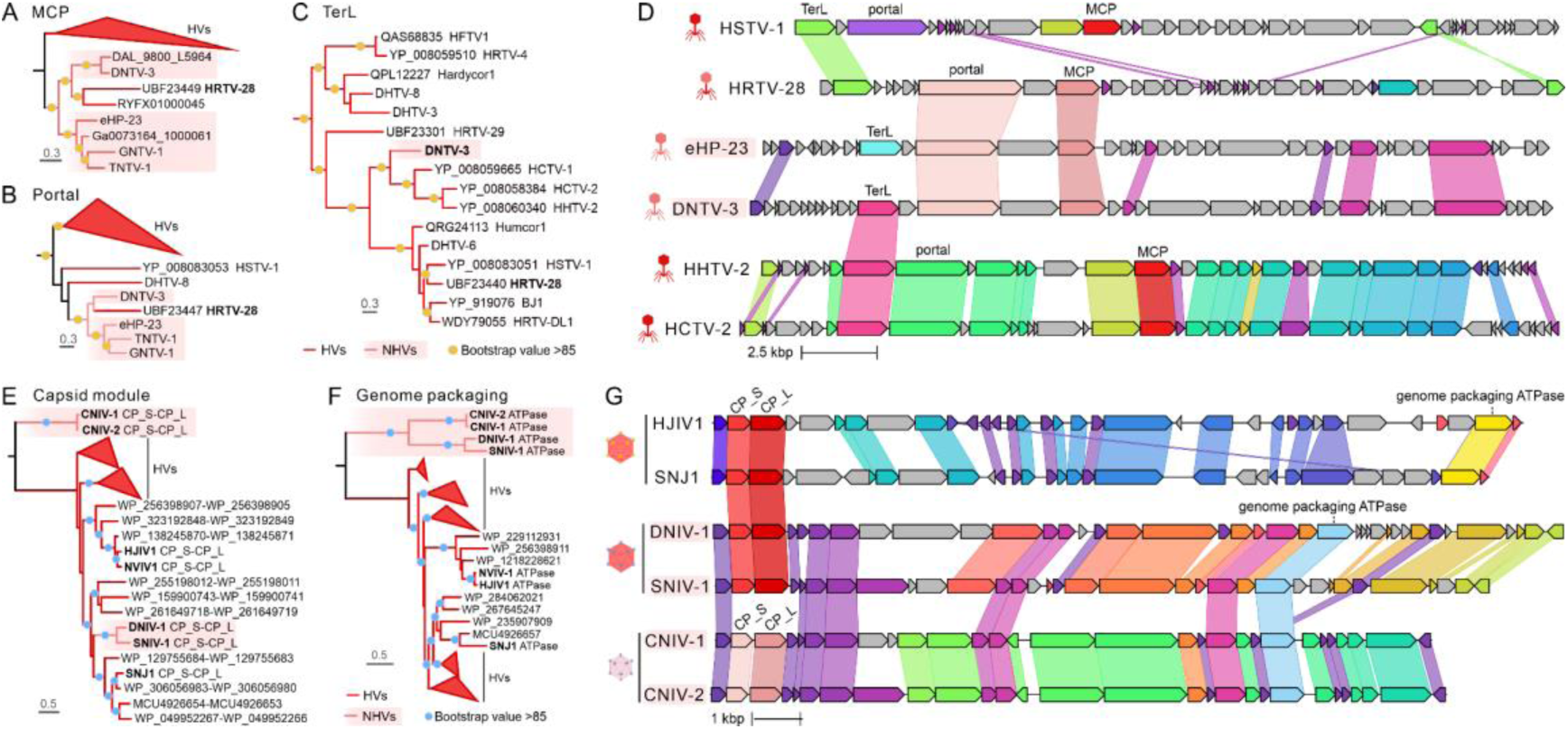
Horizontal gene transfer between HVs and NHVs. a-c. Maximum likelihood phylogenies of the major capsid proteins (MCP, panel a), large subunit of the terminase (TerL, panel b) and portal (panel c) of head-tailed HVs and NHVs. Branches with bootstrap support values higher than 85% are indicated with yellow circles. Expanded trees are shown in Supplementary figure 9. d. Comparison of the (partial) genome maps of HSTV-1 (KC117378: 1-25,344 bp), HRTV-28 (MZ334528: 1-24,686 bp), ePH23 (JQ807243: 1-13,391 bp, 18,015-30,840 bp), DNTV-3 (7,207-33,827 bp), HHTV-2 (KC292024: 12,973-38,639 bp) and HCTV-2 (KC292028: 6,250-33,408 bp). Homologous genes (>25% identity) are highlighted using the same color and linked via shadings. e-f. Maximum likelihood phylogenies of the concatenated sequences of the large and small major capsid proteins (CP_L and CP_S, respectively; panel e) and genome packaging ATPase (panel f) of tailless icosahedral HVs and NHVs. Branches with bootstrap support values higher than 85% are indicated with blue circles. g. Comparison of the genome maps of HJIV1, SNJ1, DNIV-1, SNIV-1, CNIV-1 and CNIV-2. Homologous genes (>25% identity) are highlighted using the same color and linked via shadings. Relevant genes are labaled.

The presence of more diverse AMGs in DNTV-1 genome compared to HVs with large genomes may be a reflection of the metabolic limitations of the nanohaloarchaeal host, so that viruses encoding AMGs have fitness advantage.

### Horizontal gene transfer between HVs and NHVs

Although our data suggests that HVs and NHVs did not switch hosts across the phylum boundary, blastp analysis of the viral proteomes pinpointed 18 NHV genes that appear to have been horizontally transferred between the two virus groups (Table S5). Most of these genes encode major morphogenesis proteins, such as major capsid proteins and genome packaging ATPases, and homing endonucleases. To formally assess the potential gene transfers between HVs and NHVs, we focused on the morphogenetic module genes of head-tailed and tailless icosahedral viruses. Phylogenetic analysis of the HK97-fold MCPs encoded by haloarchaeal and nanohaloarchaeal head-tailed viruses revealed that the MCP of *Halorubrum* tailed virus 28 (HRTV-28), the sole representative of the *Suolaviridae* family ^36^, was nested among the MCPs of DNTV-3-like NHVs (Fig. 5A, Fig. S10). Notably, the MCP assembly into a capsid in duplodnaviruses is nucleated by the portal protein and this functional coupling typically leads to coevolution of the two proteins ^50,51^. Indeed, results of the phylogenetic analysis of the HV and NHV portal proteins mirror those of the MCPs, with the portal protein of HRTV-28 being nested among the portal proteins of tailed NHVs (Fig. 5B, Fig. S10). These results suggest that the MCP and portal genes of HRTV-28 were simultaneously acquired by horizontal transfer from tailed NHVs. Indeed, these are the only genes of HRTV-28 that show closer similarity (∼30% identity) to the proteins of NHVs (Fig. 5D). Consistently, phylogenetic analysis of the genome packaging enzyme, the large terminase subunit (TerL), showed that the TerL of HRTV-28 clusters with homologs from other HVs. Remarkably, however, the TerL of DNTV-3 grouped with homologs from HVs rather than other NHVs, suggesting an HV-to-NHV transfer (Fig. 5C and 5D).

Exchange of the structural modules appears not to be restricted to head-tailed HVs and NHVs. The four non-tailed icosahedral NHVs share a set of nine genes and can be classified within the same new family (Fig. 2B). However, the two single jelly-roll MCPs of DNIV-1 and its close relative SNIV-1 show high sequence similarity (49.4% and 30.7% identity) with the homologs encoded by icosahedral HVs of the family *Simuloviridae* (Table S5). By contrast, the two other DNIV-1-like NHVs, CNIV-1 and CNIV-2, encode distinct MCP variants that are not recognizably similar to the simulovirus or DNIV-1 MCPs (Fig. 5G). Consistently, phylogenetic analysis placed the DNIV-1 and SNIV-1 MCPs deep within the HV clade (Fig. 5E). By contrast, the FtsK-like genome packaging ATPases of the tailless icosahedral HVs and NHVs formed distinct clades corresponding to their host associations (Fig. 5F). These results suggest that the ancestral MCPs, resembling those of CNIV-1 and CNIV-2, were replaced in DNIV-1 and SNIV-1 by homologs from HVs. Collectively, these results suggest a rather common bidirectional horizontal exchange of homologous genes between HVs and NHVs, likely facilitated by the cytoplasmic exchange between the haloarchaeal and nanohaloarchaeal hosts ^52^.

### Spindle-shaped viruses of haloarchaea and nanohaloarchaea are associated with circular satellite molecules

The signature morphogenetic module of spindle-shaped viruses includes the hydrophobic α-helical MCP and the AAA+ ATPase. Unexpectedly, assembly of the Danakil metagenomes yielded 9 complete genomes of small circular elements (2.7-9.0 kb, median 3.6 kb) that encode one or both of the two spindle-shaped virus signature proteins. Seven of these elements, hereinafter SRCEs for spindle-shaped virus-related circular elements, were targeted by haloarchaeal CRISPR spacers (Table S6). Consistently, phylogenetic analysis of the SRCE AAA+ ATPases showed that they form a monophyletic clade with the homologs encoded by spindle-shaped HVs (Fig. 6A), including DHSV-1, suggesting that theses SRCEs replicate in haloarchaea. Further searches against the IMG/VR database queried with the ATPases and MCPs of DNSVs led to the identification of four SRCE-like elements, which in the phylogenetic analysis of the AAA+ ATPases clustered with DNSV-1 and DNSV-2, suggesting nanohaloarchaeal hosts. Thus, the AAA+ phylogeny supports the early divergence of haloarchaeal and nanohaloarchaeal spindle-shaped viruses and suggests that both groups of viruses are associated with SRCEs.

**Figure 6.**
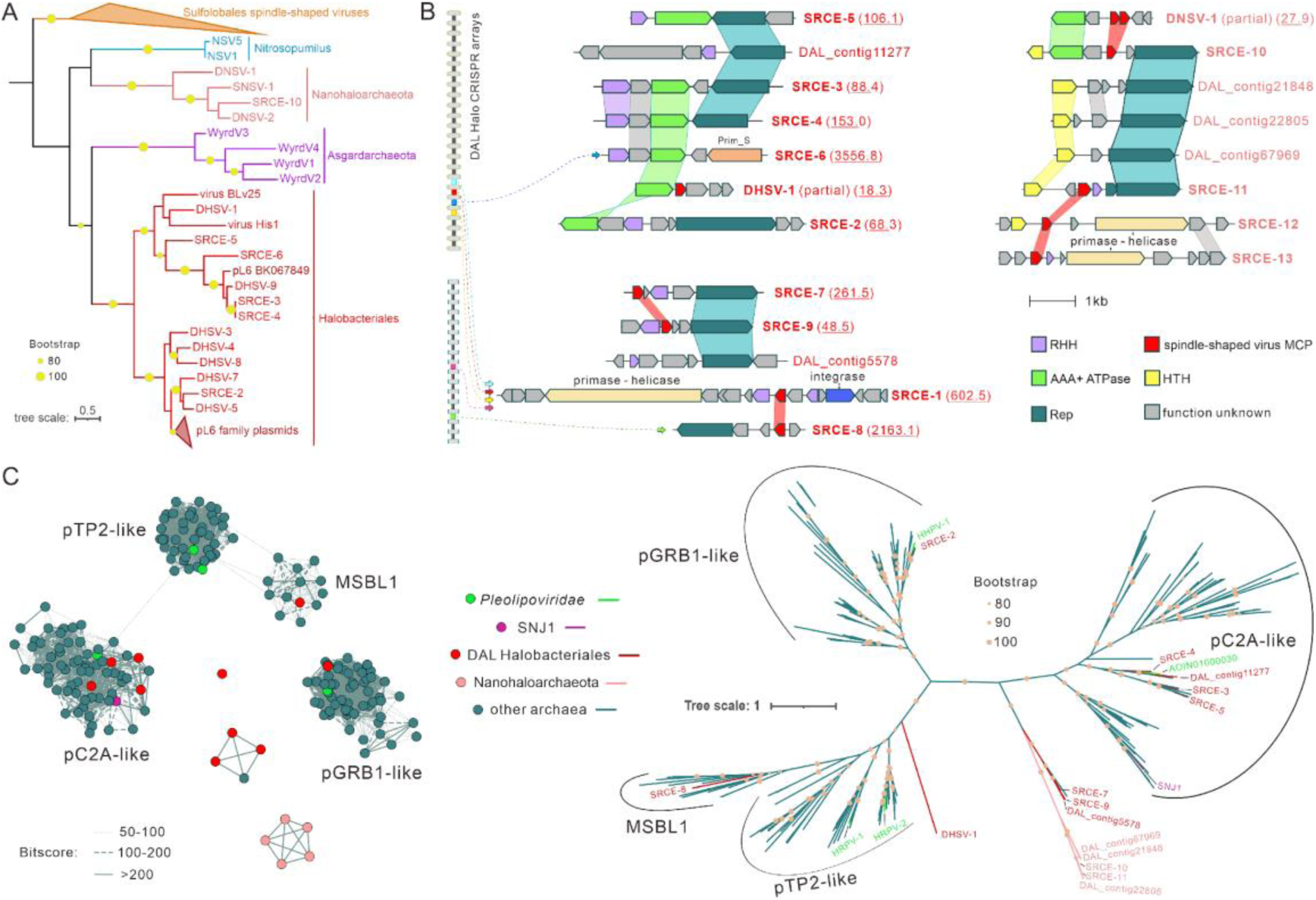
Putative satellites associated with spindle-shaped HVs and NHVs. a. Maximum likelihood phylogeny of the AAA+ ATPases conserved in spindle-shaped HVs and NHVs and SRCEs. Branches with bootstrap support values higher than 85% are indicated with yellow circles. b. Genome maps of SRCEs and related plasmids. Sequencing depth (when available) of each genome is shown in parentheses following the genome name. Functionally equivalent genes are colored using the same color. Genes with over 20% identity are linked via shadings. Spacers targeting the SRCEs are shown on the left of the figure as colored bars within the CRISPR arrays, with arrows linking the spacers with the corresponding SCREs. c. Left: similarity networks of rolling circle replication initiation endonuclease (RCRE) sequences are shown on the left. Each node represents an RCRE sequence and lines linking the nodes represent the degree of sequence similarity based on BLASTp bitscores. Right: maximum likelihood phylogeny of RCRE proteins encoded by diverse archaeal mobile genetic elements. Branches with bootstrap support values higher than 80% are indicated with pink circles.

Besides the two viral proteins, SRCEs encode plasmid-like replication proteins, namely, rolling circle replication initiation endonucleases (Reps) or archaeo-eukaryotic primases-helicases belonging to several different families (Fig. 6B and 6C). The combination of the spindle-shaped virus signature proteins with plasmid replication proteins found in SRCEs resembles the features of the satellite plasmid pSSVx of the thermoacidophilic archaeon *Saccharolobus islandicus*. Similar to SRCEs, pSSVx combines a replication module closely related to that of *Saccharolobus* plasmids with homologs of two fusellovirus SSV2 proteins, including the AAA+ ATPase^53^. pSSVx is maintained in the cells as a plasmid but upon coinfection with SSV2, pSSVx is encapsidated into smaller spindle-shaped particles and spreads horizontally in the population^53^. We hypothesize that SRCE represent satellites associated with haloarchaeal and nanohaloarchaeal spindle-shaped viruses. Notably, similar SRCE-like agents were previously documented in the hypersaline Lake Reba, Senegal^54^, but were not considered to represent virus satellites. The sequence similarity networks and the Rep phylogeny showed that SRCEs formed five different clusters, including previously described MSBL1, pGRB1-like and pC2A-like Rep families, as well as two new clusters consisting of haloarchaeal and nanohaloarchaeal SRCEs and plasmids (Fig. 6C).

By hijacking the morphogenetic module of the helper viruses, satellite nucleic acids can have adverse effects on the propagation of the coinfecting viruses^55^. We thus assessed the distribution and replicative potential of the SRCEs and spindle-shaped viruses, including those with incomplete genomes, in the Danakil metagenomes by comparing their sequencing depths (Fig. S11, Table S1, see Supplementary text). The sequencing depth of SRCEs was generally higher than that of the spindle-shaped viruses, with SRCE-6 and SRCE-8 being up to 155-fold and 94-fold more abundant than their putative helper virus DHSV-2 (3,557× and 2,163× vs 23×, respectively; see Table S1, Fig. 1G). Note that SRCE-6, SRCE-8 and DHSV-2 are targeted by the same CRISPR arrays and are all found in the Western-Canyon Lakes (Table S1, S6 and S7). Furthermore, SRCEs displayed broader distribution across Danakil lakes compared to most DHSVs (Fig. S11). Given the dominance of the SRCE genomes, it is possible that under coinfection the majority of viral structural proteins would be hijacked by SRCEs, thereby negatively impacting the production of the helper virus. Notably, seven different CRISPR arrays contained spacers against more than one SRCE (Table S7, Fig. 6B), suggesting that distinct SRCE variants may co-exist in the same host cell. Collectively, our results point to the existence of satellite agents parasitizing both haloarchaeal and nanohaloarchaeal spindle-shaped viruses, uncovering a hidden layer of parasitism in hypersaline environments.

## Discussion

Metagenomics has revolutionized the virus discovery, bringing to light the viromes associated with organisms that are difficult or impossible to cultivate under laboratory conditions. Here we applied this approach to explore the viromes associated with extreme halophiles of the phyla Halobacteriota and Nanohaloarchaeota (DPANN superphylum), which dominate microbial communities in hypersaline lakes in Danakil Depression, one of the most extreme habitats on Earth. Although genomes of several NHVs have been described previously, the extent of their diversity beyond *Caudoviricetes*, interplay with the nanohaloarchaeal hosts and evolutionary relationships to viruses of haloarchaea were not sufficiently explored. We found that both haloarchaea and nanohaloarchaea are associated with four groups of viruses, namely, head-tailed viruses (class *Caudoviricetes*, realm *Duplodnaviria*), tailless icosahedral viruses (realm *Singelaviria*), pleomorphic viruses (realm *Monodnaviria*) and spindle-shaped viruses. Notably, across the discovered virus assemblages, HVs and NHVs display comparable genome lengths, indicating that unlike their hosts, NHVs did not evolve by genome reduction. Notably, the metagenomes analyzed in this study were produced from samples retained on the 0.2-30 µm filter membranes after filtration, i.e., with the majority of extracellular virions being removed. Thus, the reconstructed viromes are likely to be enriched in viruses and satellites actively replicating within their hosts.

It is rather striking that despite the extensive phylogenetic distance between haloarchaea and nanohaloarchaea, the composition of their viromes appears to consist of the same virus groups. A priori, two scenarios are possible, (i) convergent adaptation of particular virus morphotypes to hypersaline environments and/or independently adapted lineages of halophilic archaea or (ii) host-switching between haloarchaea and nanohaloarchaea. Comparative genomics, phylogenetic analysis and comparison of the amino acid usage and nucleotide compositions strongly suggest that HVs and NHVs from all four assemblages have diverged early, likely concomitantly with the radiation of archaeal lineages, supporting the first of the two scenarios. Thus, the four virion morphotypes appear to be adapted to the hypersaline habitats. Nevertheless, we identified cases of bidirectional horizontal gene exchange between HVs and NHVs, consistent with cytoplasmic exchange, as proposed in the case of mutualistic interaction between haloarchaea and nanohaloarchaea ^11^. The exchanged genes encode for major structural and morphogenesis proteins. Notably, genes encoding the capsid protein and genome packaging enzyme are among the most preferred targets of the CRISPR systems ^56^. Furthermore, certain pattern recognition-based defense systems in bacteria and archaea are known to recognize the conserved structural and morphogenesis proteins, such as portal and terminase ^57^. We hypothesize that the observed horizontal exchange of the morphogenesis genes is, at least partly, precipitated by the need to evade different cellular defense systems.

The identified viruses represent 16 families, 12 of which were not described previously, highlighting the sparse sampling of the archaeal virosphere and the distinctiveness of the Danakil viromes. To the best of our knowledge, we describe the first spindle-shaped, pleomorphic and tailless icosahedral viruses associated with Nanohaloarchaeota. Spindle-shaped viruses are among the most widespread groups of archaeal viruses previously detected in diverse environments and hosts, and hypothesized to have infected the last archaeal common ancestor ^58,59^. Identification of such viruses in nanohaloarchaea further supports this hypothesis. The pleomorphic viruses (family *Pleolipoviridae*) were initially restricted to haloarchaeal hosts ^23,60^. However, evolutionarily related viruses were recently described in marine hyperthermophiles of the class Archaeoglobi (phylum Halobacteriota), halophilic methanogens of the class Methanonatronarchaeia (phylum Halobacteriota) and methanogenic archaea of the order Methanomassiliicoccales (phylum Thermoplasmatota) ^37,38^. The proposed new family “*Nanopleoviridae*” extends the pleomorphic virus distribution to a third phylum, Nanohaloarchaeota, suggesting a more ancient emergence of this virus lineage in archaea than previously considered. Tailless icosahedral viruses of the realm *Singelaviria* currently constitute two families, *Sphaerolipoviridae* and *Simuloviridae*, both associated with haloarchaeal hosts. We identified representatives of two additional potential *Singelaviria* families, “*Halicoviridae*” and “*Nanicoviridae*”, with the latter one extending the reach of archaeal singelaviruses to the phylum Nanohaloarchaeota. The most diverse in our dataset were viruses of the class *Caudoviricetes*, which could be assigned to 10 different families, 7 of which are new (Table S1). Two of the previously established families, namely, *Madisaviridae* and *Pyrstoviridae*, were represented by singletons and are now expanded by HVs from the Danakil viromes. Notably, none of the NHVs could be assigned to families containing HVs, emphasizing the deep divergence of viruses associated with the two groups of halophilic archaea. Consistently, none of the viral genomes was targeted by both haloarchaeal and nanohaloarchaeal spacers, suggesting that the two archaeal lineages have distinct viromes despite living in close contact. Notably, it has been recently shown that some archaea use CRISPR systems to fend against DPANN ectosymbionts ^61^. However, none of the haloarchaeal spacers retrieved in our study targeted nanohaloarchaeal genomes, which would be consistent with the possibility that nanohaloarchaea in the Danakil salt lakes act as symbionts rather than parasites of haloarchaea.

Some of the large NHVs, such as DNTV-1, carry a number of AMGs, primarily implicated in nucleotide and amino acid metabolism, and translation, and could have played an important role in the horizontal AMG transfer between viruses and cells, as in the case of the RNR. Nevertheless, it is unlikely that NHVs could complete the replication cycle in nanohaloarchaea that are not attached to the haloarchaeal cells. Thus, the dependence of nanohaloarchaea on haloarchaea for basic biomolecules, such as nucleotides and amino acids, renders NHVs secondarily dependent on haloarchaea besides their direct host nanohaloarchaea. In this context, the discovery of potential satellites, dubbed SRCEs, associated with haloarchaeal and nanohaloarchaeal spindle-shaped viruses is perhaps most striking. NHVs can be considered hyperparasites (parasites of parasites) in their own right, whereas NHV satellites add yet another layer to this nested system (Fig. S12).

Virus satellites were known in eukaryotes for decades and especially extensively studied in plants where they can impact the outcome of virus infection in a number of ways and cause pathologies ^55,62–64^. Satellites have been also known to parasitize bacterial viruses ^65^, but the full extent of their diversity, distribution and impact on bacterial communities has only recently started to emerge through both (meta)genomics and cultivation studies ^66–71^. By contrast, in the domain Archaea, only two satellites, pSSVi and pSSVx, have been described previously, both associated with spindle-shaped viruses of the family *Fuselloviridae* infecting thermoacidophilic archaea of the genus *Saccharolobus* ^53,72^. A common defining property of satellites across the cellular domains is that they are capable of hijacking the virion morphogenesis machinery of helper viruses for horizontal spread in the population. Haloarchaeal and nanohaloarchaeal SRCEs combine plasmid-derived replication modules with incomplete virion morphogenetic modules. In particular, they can encode either MCP or ATPase, or both. Notably, pSSVx associated with fuselloviruses encodes a homologous ATPase, but not the MCP. Thus, in this respect, SRCE resemble the recently discovered bacterial satellites, dubbed cf-PICI, that encode the MCPs ^70^, but appropriate the phage tails for propagation.

Although the SRCEs described herein were sequenced from the total metagenomes, we note that they are related to enigmatic elements previously detected in the virus-sized fraction of hypersaline samples filtered through 0.2 µm filter and pretreated with DNase I to remove the free DNA ^54^, suggesting that SRCEs are bona fide satellites capable of extracellular spread. Consistent with their broad distribution in hypersaline environments, we identified SRCE-like satellites in the IMG/VR database. Furthermore, SRCEs appear to be related to the globally distributed PL6-family plasmids, in which AAA+ ATPase is one of the core genes ^73,74^. Indeed, in our phylogenetic analysis, ATPases of the PL6-family plasmids were nested among those encoded by SRCEs (Fig. 6A). Thus, we propose that PL6-family plasmids represent satellites that can hijack the structural module of haloarchaeal spindle-shaped viruses upon coinfection.

The sequencing depth of SRCEs was consistently higher than that of the spindle-shaped viruses, suggesting a negative impact of the satellites on the helper viruses. According to the most straightforward (and simplistic) scenario, in the absence of virus infection, the SRCE would be maintained in the population, with low fitness cost. However, upon active virus infection, SRCE would titer out the structural proteins of the helper virus, thereby decreasing its progeny yield and limiting the spread in the population. Whether this competition benefits the host cell, with the satellite acting as a host mutualist in the tripartite consortium, remains to be addressed experimentally once the SRCE-virus-host systems are isolated. Collectively, our results underscore the complexity of natural microbial communities and provide the first glimpse at the nested hyperparasitic interactions between haloarchaea, symbiotic nanohaloarchaea, their viruses and virus satellites (Fig. S12).

## Methods

### Environmental samples

Brine samples used for this study were collected in January 2019 from different sites in the north Danakil Depression, Ethiopia: a cave reservoir in the Dallol dome salt canyons (9Gt), the Western Canyon Lakes WCL2 and WCL3, and the middle of Lake Karum or Assale close to one of its islands (9Ass)^14^. An additional sample from the northwest rim of Lake Assale was collected in 2016 ^14^. Their physicochemical parameters and hydrochemistry were characterized, with the geothermally influenced and actively degassing WCLs being the most extreme and chaotropic systems sampled in the region^14^. Microbial biomass in the range 0.2-30 µm cell diameter was fractioned by filtration, fixed in EtOH in situ and, after DNA purification in the laboratory, used for direct metagenome sequencing ^29^. In addition, unfiltered brine samples from 9Ass, 9Gt and WCL3 were collected in sterile flasks and maintained at room temperature under diel conditions for culture enrichment attempts.

### Enrichment cultures and VLP observation

Enrichment cultures of the brine samples were established in CA medium ^75^, by inoculating a 10 ml aliquot of the brine sample into 40 ml medium. The mixtures were incubated aerobically at 37°C and 100 rpm for two weeks. Cell-free culture supernatants containing VLPs were collected by centrifugation (10,000 rpm, 30 min, Beckman JLA 16.250 rotor), and concentrated by ultracentrifugation (40,000 rpm, 3h, Beckman rotor 70 Ti). The pellets were suspended in the sample buffer (2.46 M NaCl, 85 mM MgSO_4_·7H_2_O, 88 mM MgCl_2_·6H_2_O, 56 mM KCl, 3 mM CaCl_2_, 12 mM Tris-HCl (pH 7.5)). For electron microscopy observations, samples were applied to carbon-coated copper grids, negatively stained with 2% uranyl acetate and observed under the transmission electron microscope FEI Tecnai Biotwin.

### Identification of viral sequences

The five Danakil metagenomes analyzed in this study were reported recently ^29^ and are available in GenBank under BioProject PRJNA541281. The accession numbers for individual metagenomes are SAMN37693137 (DAL-Ass), SAMN37693138 (DAL-9Ass), SAMN37693139 (DAL-9Gt), SAMN37693140 (DAL-WCL2) and SAMN37693141 (DAL-WCL3). The general workflow of identification of archaeal viruses in metagenomes has been recently described in detail elsewhere ^76^. Briefly, putative viral sequences were sorted out from Danakil metagenome assemblies using two tools, geNomad (github.com/apcamargo/genomad/)^32^ and VirSorter2 (github.com/jiarong/VirSorter2)^33^. Sequences with virus score over 0.70 (geNomad or VirSorter2) were selected and potential host contaminations were removed using CheckV (bitbucket.org/berkeleylab/checkv)^77^.

### Construction of virus gene-sharing networks

Virus gene-sharing networks were built by using vConTACT v.2.0 with default parameters^78^. Input sequences were Danakil viral sequences (≥5 kb) and archaeal viral sequences and bacteriophage sequences downloaded from NCBI Reference Sequence Database (RefSeq release 220). The network file produced by vConTACT v.2.0 were visualized by Cytoscape v3.10.1 ^79^.

### Taxon-specific CRISPR identification and spacer extraction

Genomes of representative species of Halobacteriota and Nanohaloarchaeota were downloaded from Genome Taxonomy Database (GTDB, release 08-RS214) and CRISPR sequences were extracted from the genomes using CRISPRCasFinder ^80^. Redundancy removal of CRISPR sequences was done by CD-HIT clustering^81^ with a 100% sequence identity cut-off. Next, in order to get taxon-specific CRISPRs, for example, Halobacteriota-specific CRISPRs, the following procedure was conducted: all CRISPR sequences extracted from Halobacteriota were used as queries to perform BLASTn searches against genomes of all other archaeal phyla within the GTDB. CRISPR sequences were removed if they showed significant matches (>90% identity and >90% coverage) to genomes from other archaeal phyla, thus retaining only those that are unique to Halobacteriota (i.e., Halobacteriota-specific CRISPRs). The same process was applied to obtain Nanohaloarchaeota-specific CRISPRs. In total, 988 non-redundant Halobacteriota-specific and 4 non-redundant Nanohaloarchaeota-specific CRISPR sequences were obtained from the Genome Taxonomy Database (GTDB) database. Using these taxon-specific CRISPRs as references, the presence of Halobacteriota and Nanohaloarchaeota CRISPR arrays within Danakil metagenomes were identified using BLASTn search with the parameters: -word_size 7, -dust no, -perc_identity 90, - qcov_hsp_perc 90. Next, spacers were retrieved from these taxon-linked CRISPR arrays using CRISPRCasFinder. Additionally, to augment the spacer dataset for the subsequent establishment of virus-host connections, we extracted Halobacteriota and Nanohaloarchaeota CRISPR spacers also from the Earth’s microbiomes (EM) dataset using the same approach.

### Spacer targeting analysis

To make linkages between Danakil viruses and archaeal host groups, Halobacteriota and Nanohaloarchaeota CRISPR spacers were used as queries to search against viral sequences using BLASTn with the parameters: -word_size 7, -dust no ^82,83^. Consequently, a viral sequence hit showing ≥30 bp exact match with a certain spacer was considered as the spacer target (i.e., protospacer), and the spacer-related archaeal taxon was then assigned as the host group to the corresponding protospacer-carrying viral sequence. If some members of a group of closely related viruses were targeted by CRISPR spacers, all members of that group were considered to infect the same hosts. For instance, DNSV-1 is not targeted by Nanohaloarchaeota CRISPR spacers, but its two close relatives, DNSV-2 (near-complete genome) and SNSV-1 (complete genome) are both matched by Nanohaloarchaeota spacers (Fig. S3A). Thus, all members of this group were considered to infect nanohaloarchaeal hosts. Notably, DNTV-1 was found to be closely related to a viral fragment found in a Nanohaloarchaeota single-cell amplified genome (Fig. S4A)^27^.

### Viral sequence extension

The de novo assembled viral sequences (≥3 kb) were pooled together and used as seed sequences for reference assembly as previously described ^84^, in an attempt to get complete genomes. Briefly, all Danakil metagenomic sequencing reads were mapped to seed sequences using Geneious Prime with parameters: 35 bp overlap, 97% identity. The final extended sequences with ≥50 bp direct or inverted terminal repeats were considered as complete genomes.

### Virus genome annotation and phylogenomic analysis

Open reading frames (ORFs) of viral sequences were predicted using Prokka ^85^. Functional annotation was performed using HHsearch ^86^ against Pfam, PDB, CDD and viral protein sequence (uniprot_sprot_vir70) databases available from the MPI Bioinformatics Toolkit ^87^. The proteomic trees were generated using the ViPTree server version 4.0 ^88^.

### Identification of relatives of Danakil Nanohaloarchaeota viruses

Sequences of major capsid proteins of Danakil Nanohaloarchaeota head-tailed viruses, major capsid proteins and genome-packaging ATPases of Danakil Nanohaloarchaeota tailless icosahedral viruses, pleolipovirus VP4-like proteins and AAA+ ATPases of Danakil Nanohaloarchaeota pleomorphic viruses, and major capsid proteins and AAA+ ATPases of Danakil Nanohaloarchaeota spindle-shaped viruses were used as queries to perform BLASTp searches in NCBI nr database and IMG/VR database. Hits with ≥30% identity, ≥50% coverage were retrieved and the corresponding viral sequences were marked as relatives of Danakil Nanohaloarchaeota viruses and downloaded for following analyses.

### Orthologous fraction calculation of the viral genomes

The proportion of orthologous fraction (OF) between the viral genomes (same inputs as for ViPTree) was estimated as previously described^76^.

### Distribution of HVs and NHVs across Danakil salt lakes

The abundance of the viral and SRCE genomes across the different metagenomes was calculated with CoverM v0.6.1 ^89^ by mapping the reads from the 5 Danakil metagenomes to each of the genome contigs. The trimmed mean was selected for visualization in a heatmap via an *ad hoc* R script using the gplots package. The dendrograms were computed and reordered based on the means. In Fig 1G, manual breaks of the data were set for min (0), median (0.6043255), mean (109.1317), and max (3369.535) for better visualization, whereas in Fig. S11, manual breaks of the data were set for min (0), median (1.845191), mean (85.48989), and max (2669.926).

### Phylogenetic analyses

a. Head-tailed haloarchaeal viruses. The major capsid protein sequences of Danakil head-tailed NHVs and HVs were pooled with their homologues from all cultured haloarchaeal viruses. Protein sequences were first clustered using CD-HIT with a 75% identity threshold (option “-c 0.75”)^81^. Next, the non-redundant sequences were aligned using Muscle5 with default parameters^90^, and non-informative columns with were removed from the alignment using trimAl v1.2 with option -gt 0.2 ^91^. Next, a phylogenetic tree was constructed based on the trimmed alignment using IQ-TREE v2.2.2.2 with the following parameters: -m MFP, -alrt 1000 ^92^. The resulting phylogeny was visualized using the iTOL online tool^93^. The same process was applied to generate phylogenetic trees for portal proteins and terminase large subunits of head-tailed haloarchaeal viruses. The trimmed alignments for the MCP, portal and TerL sequences included 452, 889 and 537 positions, and the best fitting models for phylogenetic reconstructions were Blosum62+F+I+R3, Q.pfam+F+I+G4 and Q.pfam+F+R5, respectively.

b. Tailless icosahedral NHVs. Phylogenetic trees were constructed for capsid proteins and genome packaging ATPases. The homologs of NHVs’ small capsid proteins, large capsid proteins and genome packaging ATPases were collected from the NCBI nr database through BLASTp searches (>30% identity and >50% coverage). The methods for sequence redundancy removal, sequence alignment and trimming, and tree construction and visualization were the same as employed for generating the MCP tree of head-tailed haloarchaeal viruses. Prior to tree construction, the trimmed sequence alignments of small and large capsid proteins were concatenated. The trimmed alignments of the CP_S-CP_L and genome packaging ATPase included 423 and 269 positions, respectively. The best fitting model for the CP_S-CP_L and ATPase phylogenies were Q.pfam+F+I+R4 and Q.pfam+I+G4, respectively.

c. Spindle-shaped viruses and their satellites. Reference sequences of AAA+ ATPases were extracted from genomes of spindle-shaped viruses of Sulfolobales (n=57), Asgardarchaeota (n=4), *Nitrosopumilus* (n=2), Halobacteria (n=2). The phylogenetic tree was generated using the same methods as detailed in a. The trimmed alignment included 198 positions and the best fitting model was LG+I+R4.

d. Ribonucleotide reductase (RNR) large subunits of large head-tailed NHVs

The homologs of NHV RNR large subunits were collected from NCBI Protein Reference Sequences database (Archaea, Bacteria, and Viruses) and Danakil metagenomes through BLASTp searches (>50% identity and >75% coverage). Protein sequences were initially clustered using CD-HIT with a 75% identity threshold^81^. Subsequently, the RNR large subunits of NHVs were aligned with their non-redundant homologs using Muscle5 with default parameters^90^. Non-informative columns were eliminated from the alignment using trimAl v1.2 with the option “-gappyout” ^91^. Finally, a maximum likelihood phylogenetic tree was calculated with IQ-TREE v2.2.2.2 with the parameters: -m MFP, - alrt 1000. The trimmed alignment included 477 positions and the best fitting model was Q.pfam+I+R10.

e. Glucosamine 6-phosphate synthase (GlmS) of DNTV-1

The homologs of DNTV-1 GlmS were collected from NCBI Protein Reference Sequences database (Archaea, Bacteria, and Viruses) and DAL metagenomes through 3 iterations of PSI-BLASTp searches. The phylogenetic tree was generated using the same methods as detailed in a. The trimmed alignment included 605 positions and the best fitting model was LG+F+R10.

f. Rolling circle replication initiation endonucleases (Reps) of SRCEs

The homologs of SRCE Reps were collected from ^94^ (only archaeal and archaeal virus homologs were kept). The phylogenetic tree was generated using the same methods as detailed in a. The trimmed alignment included 628 positions and the best fitting model was VT+F+R8.

## DATA AVAILABILITY

All assembled genomes were deposited to GenBank (viruses: PQ827550-PQ827567; SRCEs and plasmids: PQ766422-PQ766435).

## ACKNOWLEDGEMENTS

This work was supported by grants from Ville de Paris (Emergence(s) project MEMREMA) and Agence Nationale de la Recherche (ANR-23-CE13-022 and ANR-21-CE11-0001) to M.K., and the Moore Foundation (https://doi.org/10.37807/GBMF9739), the ANR (ANR-23-CE02-0016-01) and the European Research Council (ERC-2023-AdG 101141745) to P.L.-G. The authors thank Xue Wang, Tianqi Xu, and Anqi Zhou for their help with configuration of software and scripts.

## SUPPLEMENTARY TEXT

### AMGs encoded by HVs and NHVs

For genome replication, these viruses encode only key replisome components that are expected to recruit the rest of the host replisome through protein-protein interactions. For instance, DHTV-3, -5 and -8 only encode the PCNA sliding clamp, DHTV-4 encodes a homolog of the host replication initiator Orc1/Cdc6, DHTV-7 encodes the MCM helicase, whereas DNTV-2 encodes a homolog of the ssDNA-binding protein RPA1 (Fig. 4 and Table S3). The auxiliary genes encode proteins primarily involved in counterdefense, including diverse DNA methyltransferases, MazG-like nucleotide pyrophosphohydrolase, which is thought to prevent the programmed cell death by degrading the intracellular ppGpp ^1^, and queuine/archaeosine tRNA-guanine transglycosylase implicated in DNA modification.

The two DHTVs with the largest genomes, DHTV-1 (109.7 kb) and DHTV-2 (109.5 kb), encode nearly complete replisomes, including replicative MCM helicase, DNA primase, family B DNA polymerase, DNA polymerase sliding clamp (PCNA), sliding clamp loader (replication factor C) and DNA ligase (Fig. 4 and Table S3). Similar observations were made for large (∼100 kb) tailed viruses infecting halophilic ^2^ and marine ^3,4^ archaea, suggesting that large archaeal viruses have the capacity to semi-autonomously replicate their genomes.

DHTV-1 and DHTV-2 encode a number of proteins implicated in DNA modification and nucleotide metabolism. DHTV-1 encodes a thymidylate synthase, whereas DHTV-2 encodes a thymidylate kinase, inosine-5’-monophosphate dehydrogenase and nucleoside 2-deoxyribosyltransferase, and both viruses carry genes for dCTP deaminases (Fig. 4 and Table S3). In addition, DHTV-1 and DHTV-2 carry genes for 7-deazaguanine synthase and multiple DNA methyltransferases, which could confer resistance to type II restriction modification systems. Overall, these results are consistent with the previous notion that the most common AMGs in the HV genomes are associated with nucleotide metabolism, with genes encoding enzymes involved in other metabolic pathways being uncommon ^2^.

Besides, the proteins implicated in nucleotide metabolism and translation (Fig. 4 and Table S3), DNTV-1 encodes two sulfatase (ORF68 and ORF120), two ATP-grasp (ORF22 and ORF64) and an alkaline phosphatase (ORF146) superfamily enzymes for which substrates cannot be confidently predicted. In addition, ORF121, ORF122 and ORF135 encode glycosyltransferases, whereas ORF130 encodes a galactose-binding lectin.

### Evolutionary history of ribonucleoside-diphosphate reductases encoded by NHVs

The DNTV-1 RNR formed a clade with homologs from nanohaloarchaea and this clade was nested among RNR of haloarchaea and HVs (Fig. 5A). The topology is consistent with a scenario under which nanohaloarchaea recruited the *RNR* gene from their haloarchaeal hosts, and this gene was subsequently horizontally acquired by DNTV-1. The RNR of LTV2 was nested within a distinct small clade comprising nanohaloarchaeal homologs and lodged deep inside a clade of RNRs encoded by bacteriophages. Notably, at the base of the nanohaloarchaeal subclade was a homolog encoded by the halophilic bacterium *Salinibacter ruber* and one by the halophilic archaeon *Halorubrum ezzemoulense*. These observations suggest that DNTV-1 and LTV2 gained their respective *RNR* genes by similar but independent routes.

### Distribution of the spindle-shaped viruses and SRCEs across the Danakil metagenomes

The complete spindle-shaped viral genomes (DHSV-1, DNSV-1, and DNSV-2) appear to be restricted to specific lakes, following an opposite trend to the widespread SRCEs. We hypothesized that this might represent a snapshot of population dynamics in which the SRCEs are blooming at the expense of the fitness of their helper viruses, resulting in the depletion of the latter from certain sampling sites. To investigate this possibility, we mapped the Danakil metagenome reads against partial DHSV genomes (DHSV-2 to DHSV-11) and visualized them alongside the complete genomes of spindle-shaped viruses and SRCEs (Figure S11). The partial DHSV genomes were found to be present at similar abundance levels as the complete DHSVs. Although most of the partial DHSV genomes followed the same restriction trend observed for the complete genomes, two— DHSV-5 and DHSV-9—were widely distributed across various lakes. This could reflect the dynamics occurring when SRCEs are replicating, driving the DHSV abundance to very low levels. This scenario adds another layer of complexity to the ever-changing environment, which is influenced by the irregular input of gases and organic materials from hydrothermal activity. These fluctuations prompt the haloarchaeal community to alternate between famine and feast strategies^5^, potentially providing an opportunity for DHSVs and nanohaloarchaea to thrive opportunistically by exploiting haloarchaea. Consequently, the SRCEs, benefiting from the DHSVs, could replicate and become abundant, thereby diminishing the DHSV population.

## SUPPLEMENTARY FIGURES

**Figure S1.**
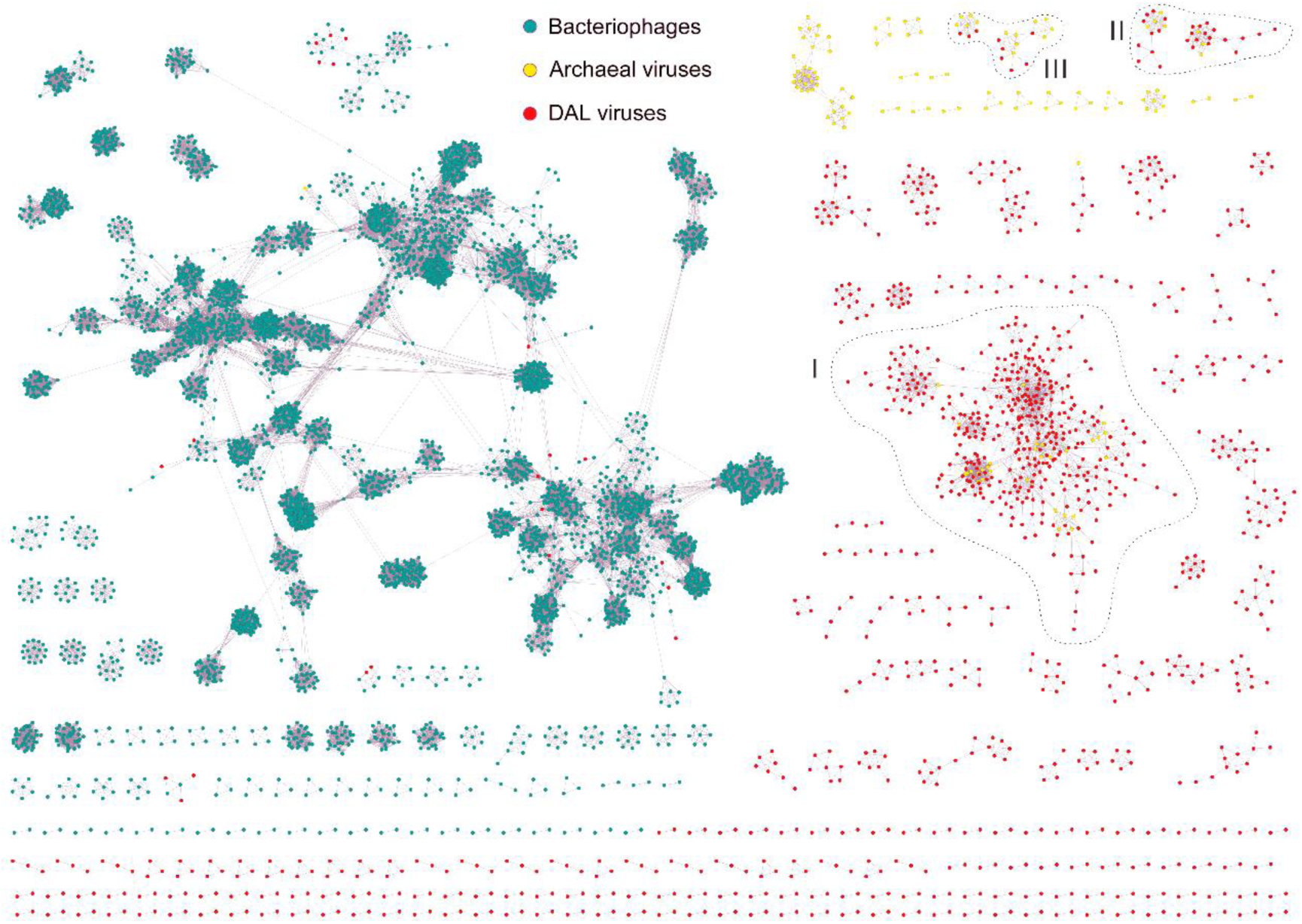
The vConTACT2 gene-sharing networks of viruses from Danakil Depression and reference prokaryotic DNA viruses. Each node represents a viral sequence and the edges between nodes represent the degree of connectivity based on the fraction of shared proteins. Nodes for reference bacteriophages are colored cyan, reference archaeal viruses are in yellow, and Danakil viruses are in red. The Danakil viral contigs formed three clusters (outlined) with previously described archaeal viruses: I, head-tailed HVs (class *Caudoviricetes*, n=382); II, tailless icosahedral HVs (families *Simuloviridae* and *Sphaerolipoviridae*, n=26); III, pleomorphic HVs (family *Pleolipoviridae*, n=9).

**Figure S2.**
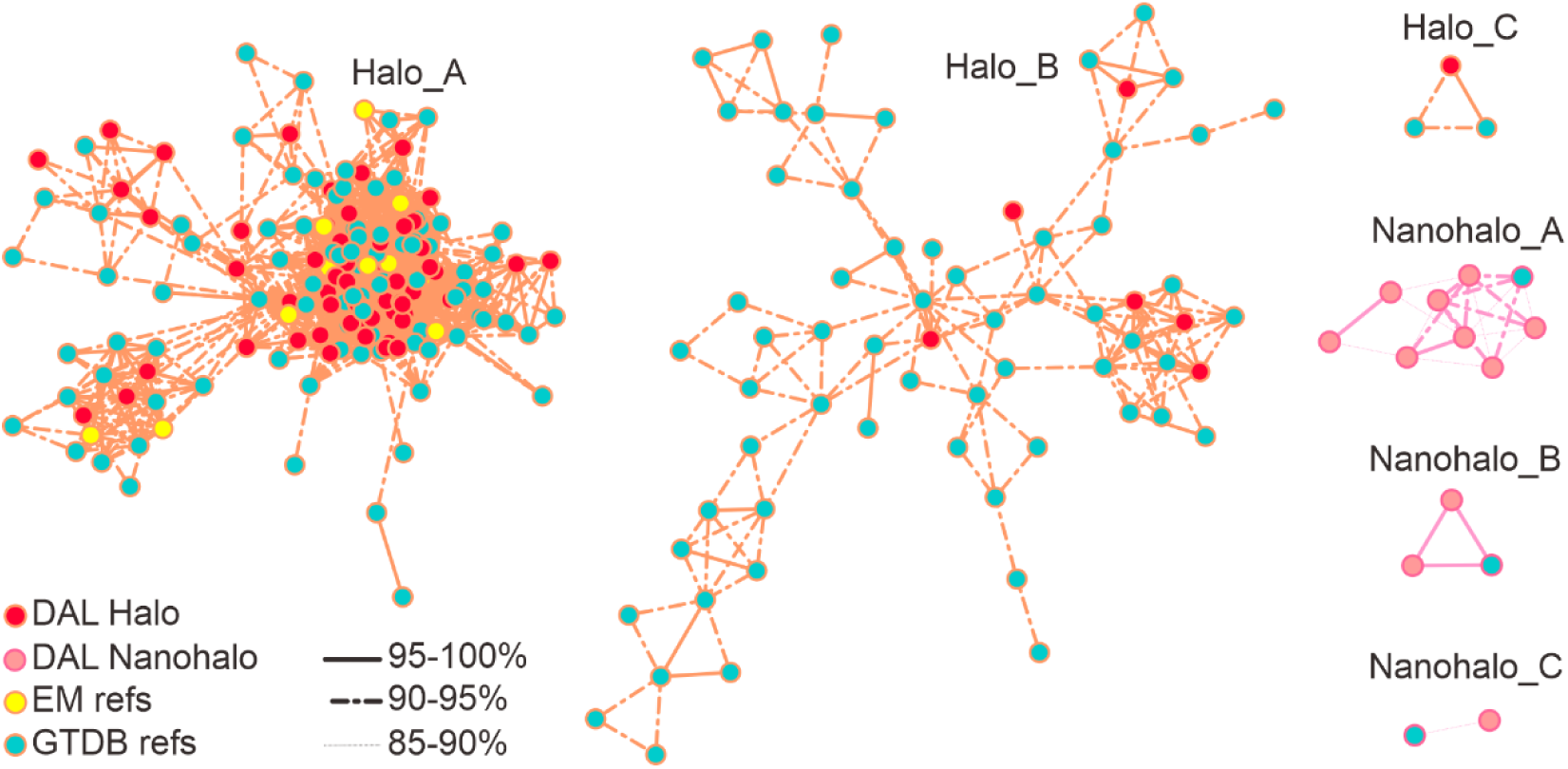
CRISPR diversity in the genomes of Halobacteriota (Halo) and Nanohaloarchaeota (Nanohalo). Similarity networks of CRISPR sequences. Each node represents a CRISPR sequence and the links between nodes represent the degree of sequence similarity between CRISPR sequences.

**Figure S3.**
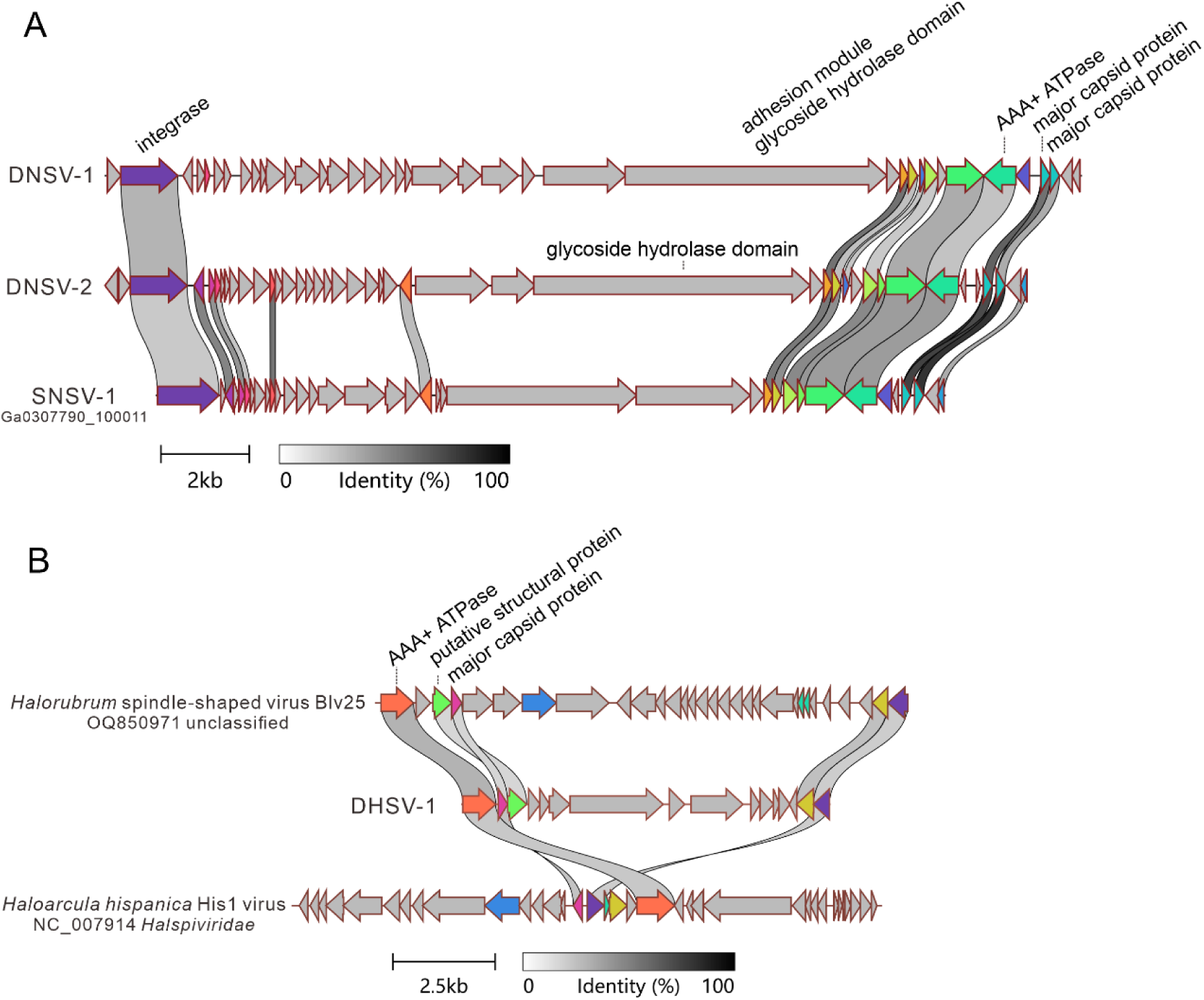
Genome maps showing the relationship among spindle-shaped viruses. a. Spindle-shaped NHVs. b. Spindle-shaped HVs.

**Figure S4.**
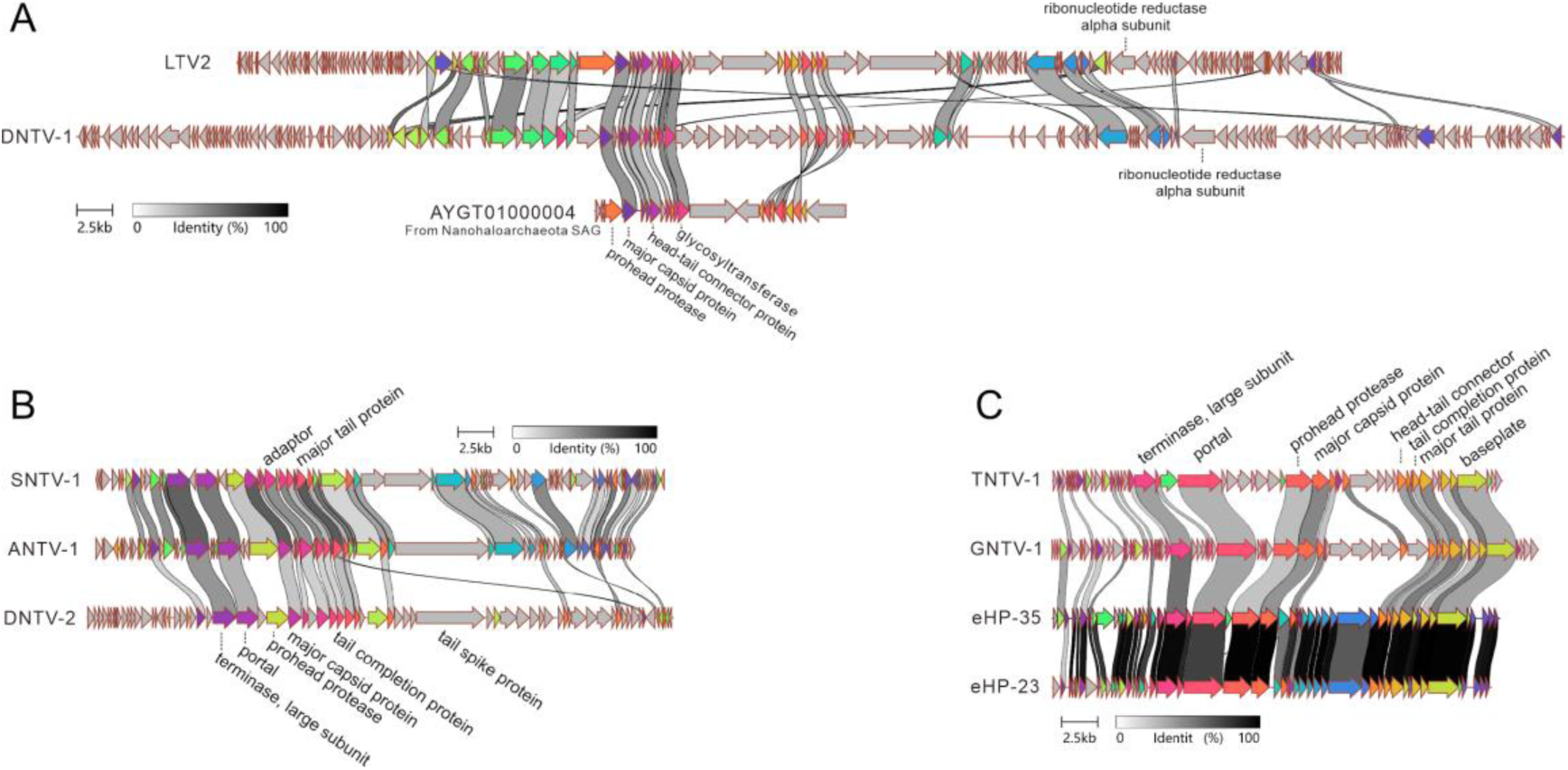
Genome maps showing the relationships among head-tailed NHVs. a. *Gulliviridae*. b. *Lilliviridae*. c. *Saladoviridae*.

**Figure S5.**
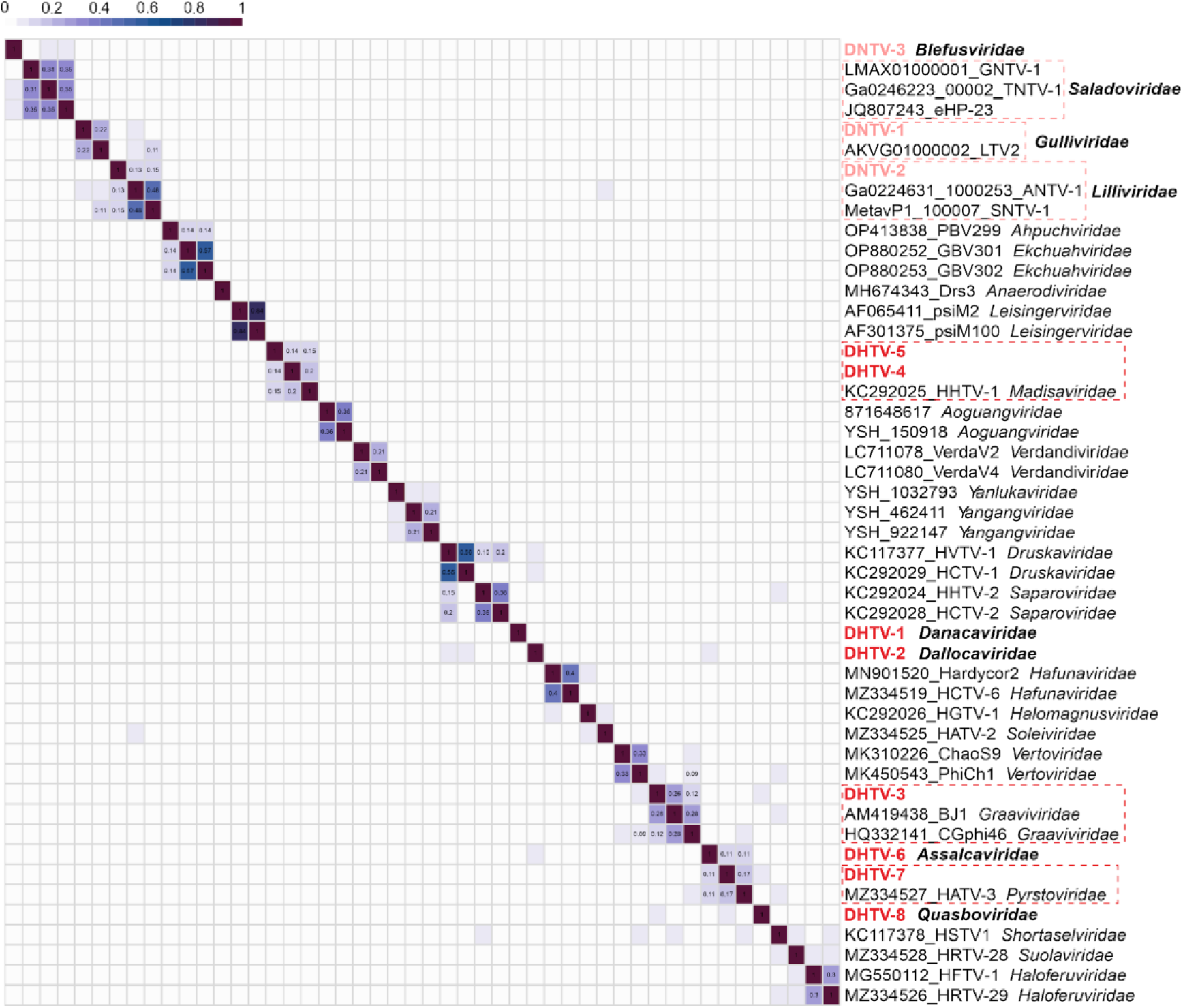
The heatmap of orthologous fraction among archaeal tailed viruses. HVs and NHVs described in this work are shown in red. Family-level groups (values >0.08) of viruses including representatives from the Danakil Depression are boxed.

**Figure S6.**
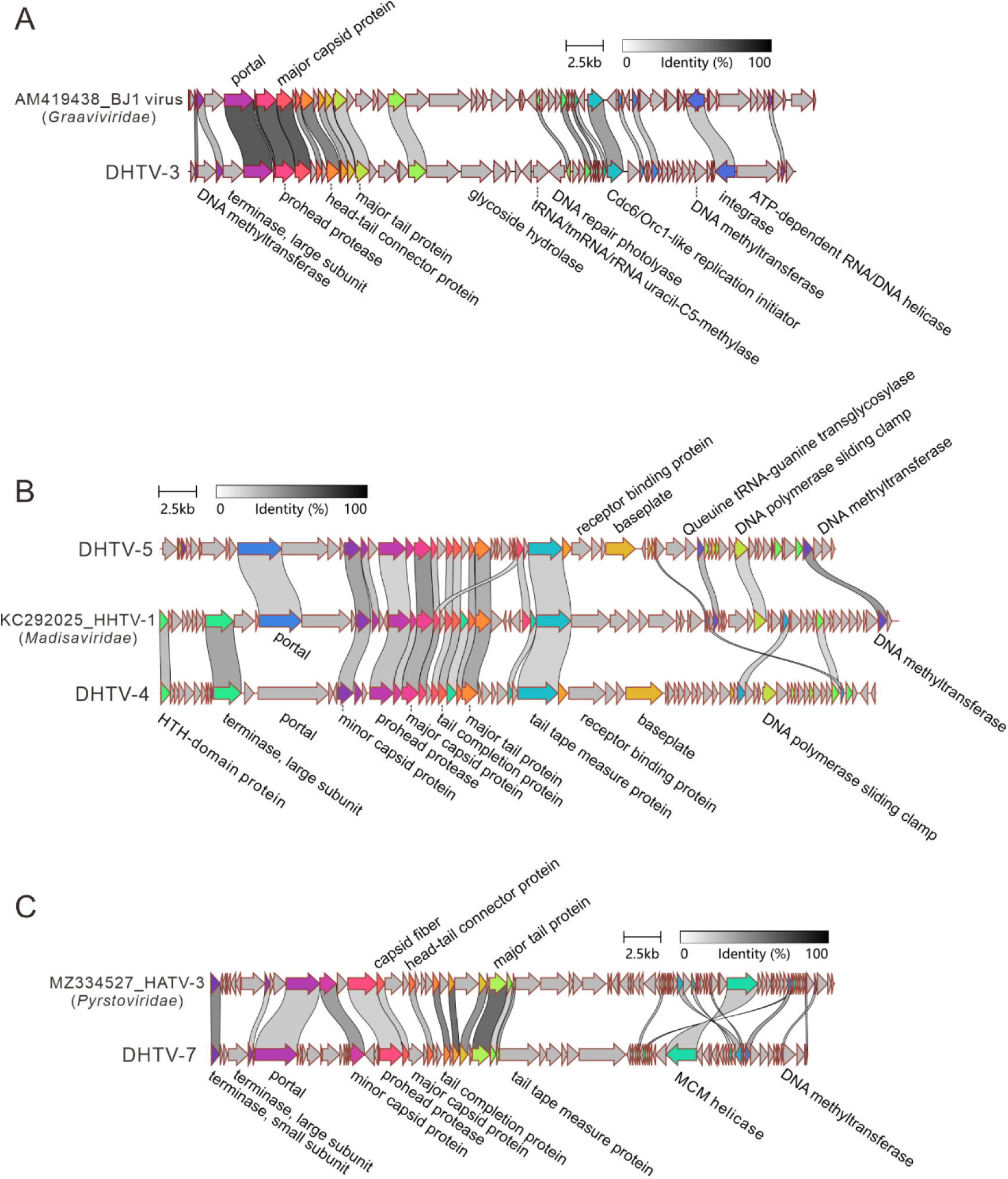
Genome maps showing the relativeness among head-tailed HVs: (A) *Graaviviridae*; (B) *Madisaviridae*; (C) *Pyrstoviridae*.

**Figure S7.**
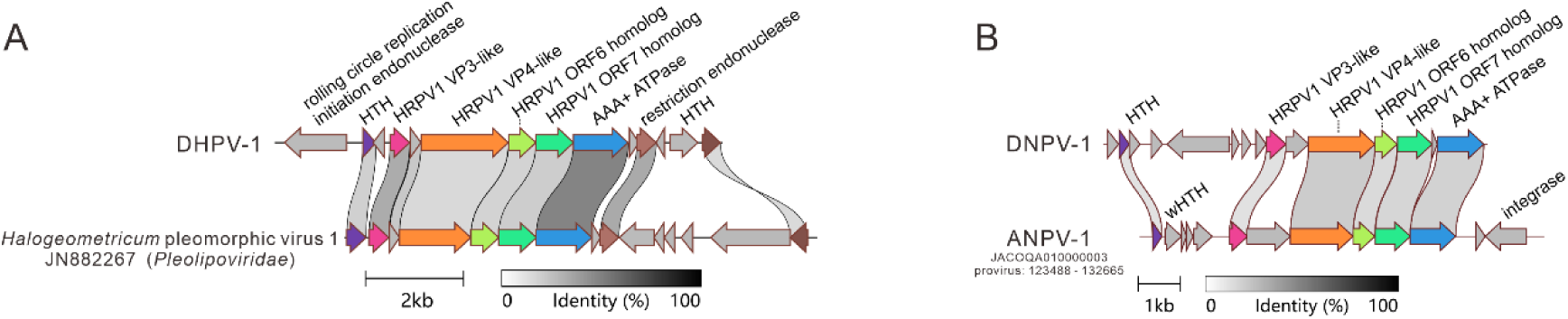
Genome maps showing the relationships among pleomorphic viruses. a. *Pleolipoviridae* HVs. b. *Nanopleoviridae* NHVs.

**Figure S8.**
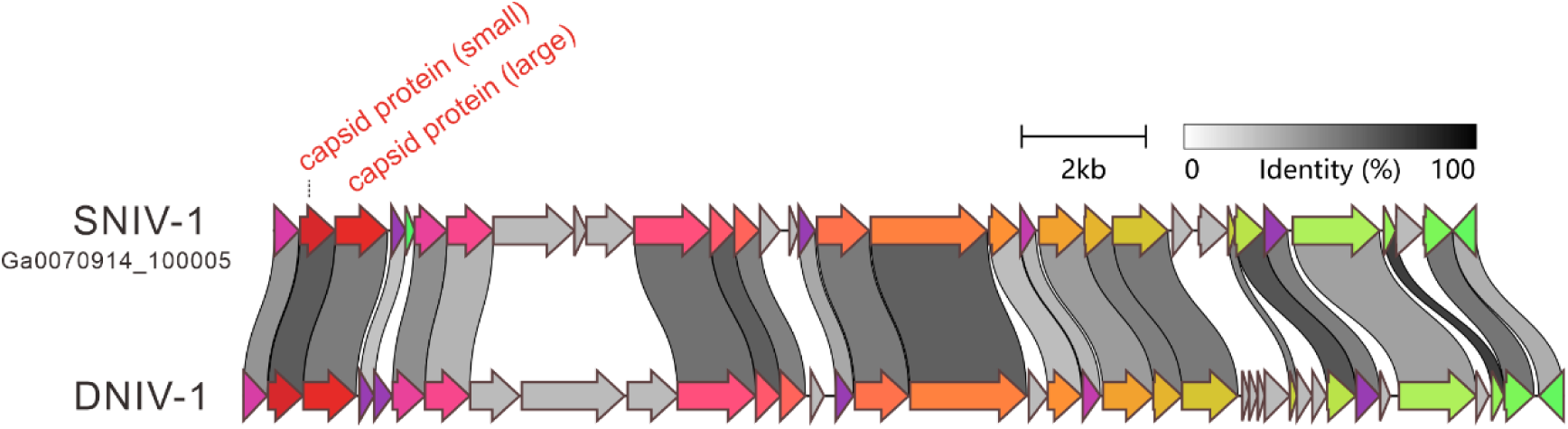
Genome maps showing the relationship among tailless icosahedral NHVs (*Nanicoviridae*).

**Figure S9.**
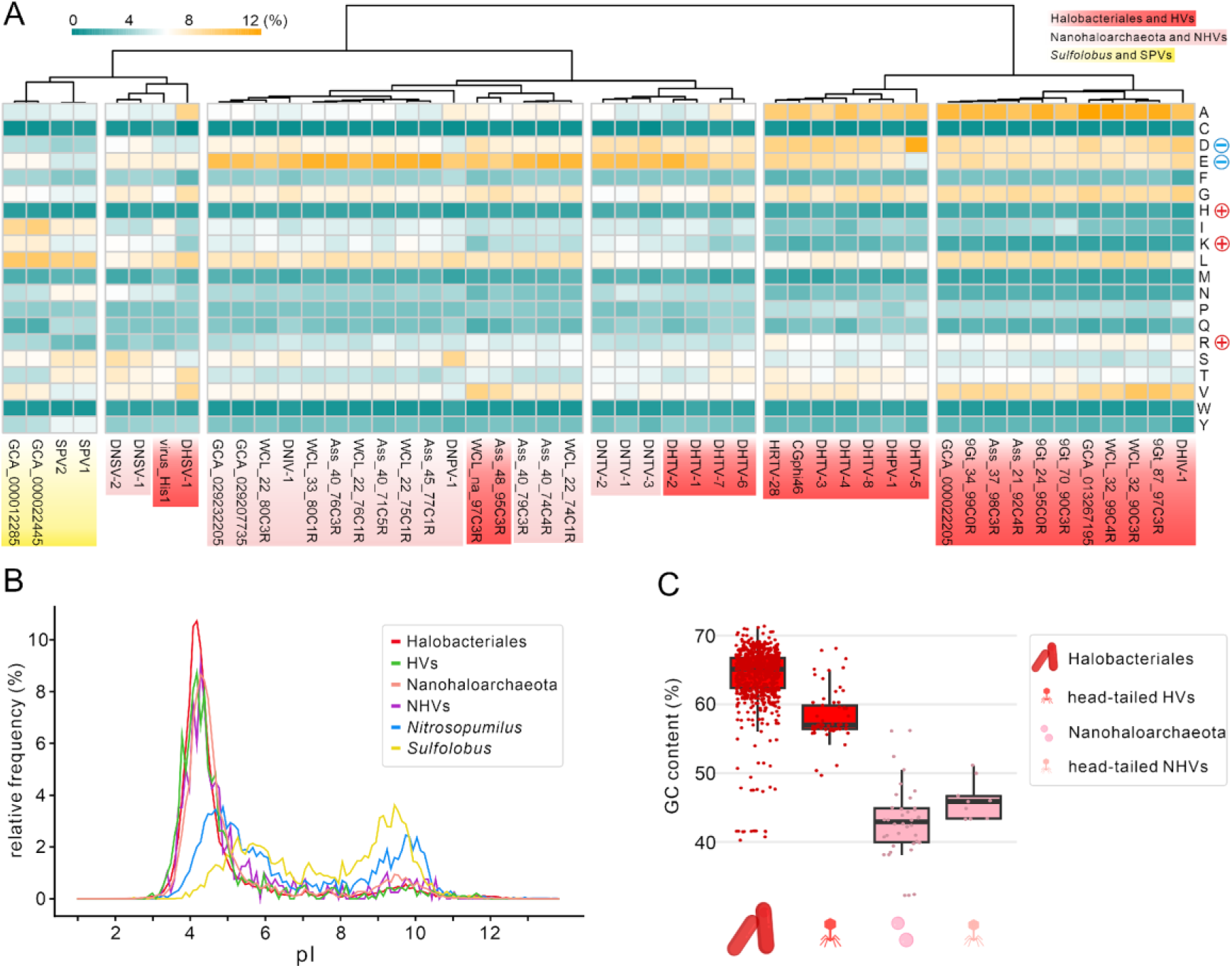
Adaptation of HVs and NHVs to hypersaline environments. a. Heatmap of amino acid usage patterns. Amino acid frequencies were calculated using proteins encoded by haloarchaea (n=10), nanohaloarchaea (n=10) and their viruses (HVs and NHVs) from the Danakil Depression. Genomes of two head-tailed HVs (CGphi46 and HRTV-28) and their hosts as well as two complete genomes of nanohaloarchaea were used for comparison. Genomes of two *Sulfolobus* species and their viruses SPV1 and SPV2 were also used for comparison. b. Distribution of isoelectric point (pI) values inferred for proteins encoded by the Danakil haloarchaea, nanohaloarchaea and their viruses (HVs and NHVs) in comparison with representative archaeal genomes from seawater (Nitrosopumilus, n=2) and hot springs (*Sulfolobus*, n=2). c. GC content (%) of genomes of Halobacteriales (n=749), Nanohaloarchaeota (n=49) and head-tailed viruses (HVs, n=62 and NHVs, n=9).

**Figure S10.**
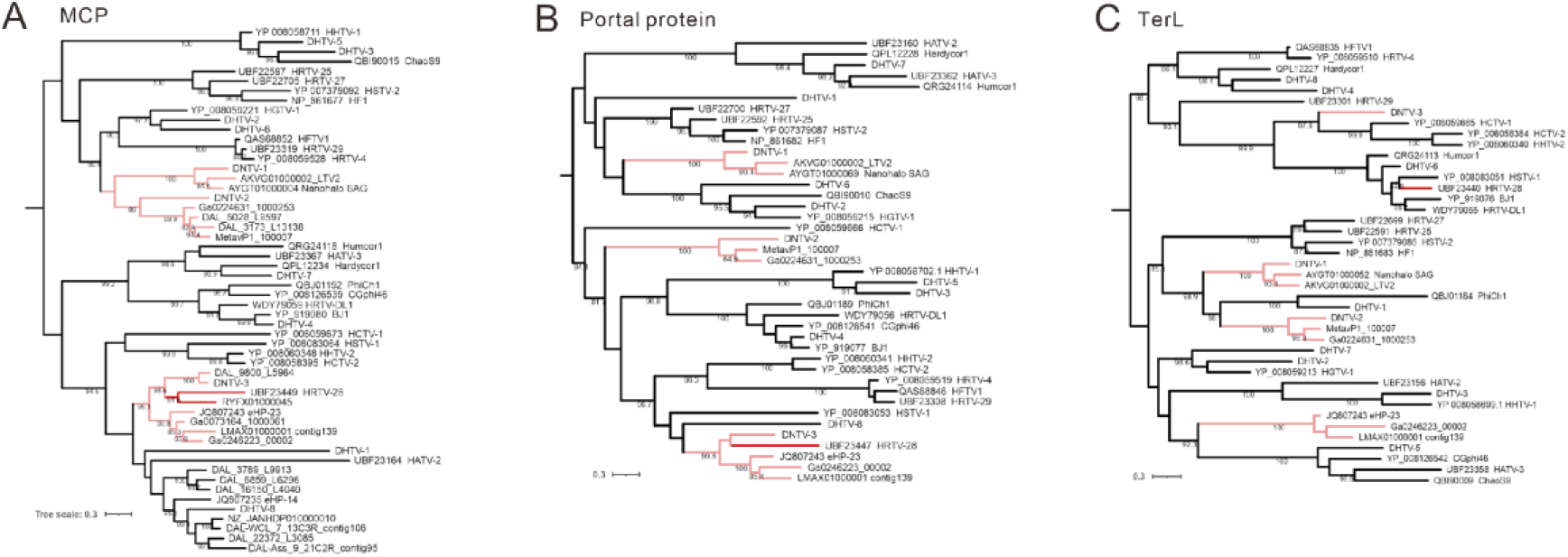
Maximum likelihood phylogenies of the hallmark proteins of head-tailed viruses. a. Major capsid protein (MCP) of head-tailed HVs and NHVs. b. Portal protein of head-tailed HVs and NHVs. c. Terminase large subunit (TerL) of head-tailed HVs and NHVs. NHVs are indicated with pink branches.

**Figure S11.**
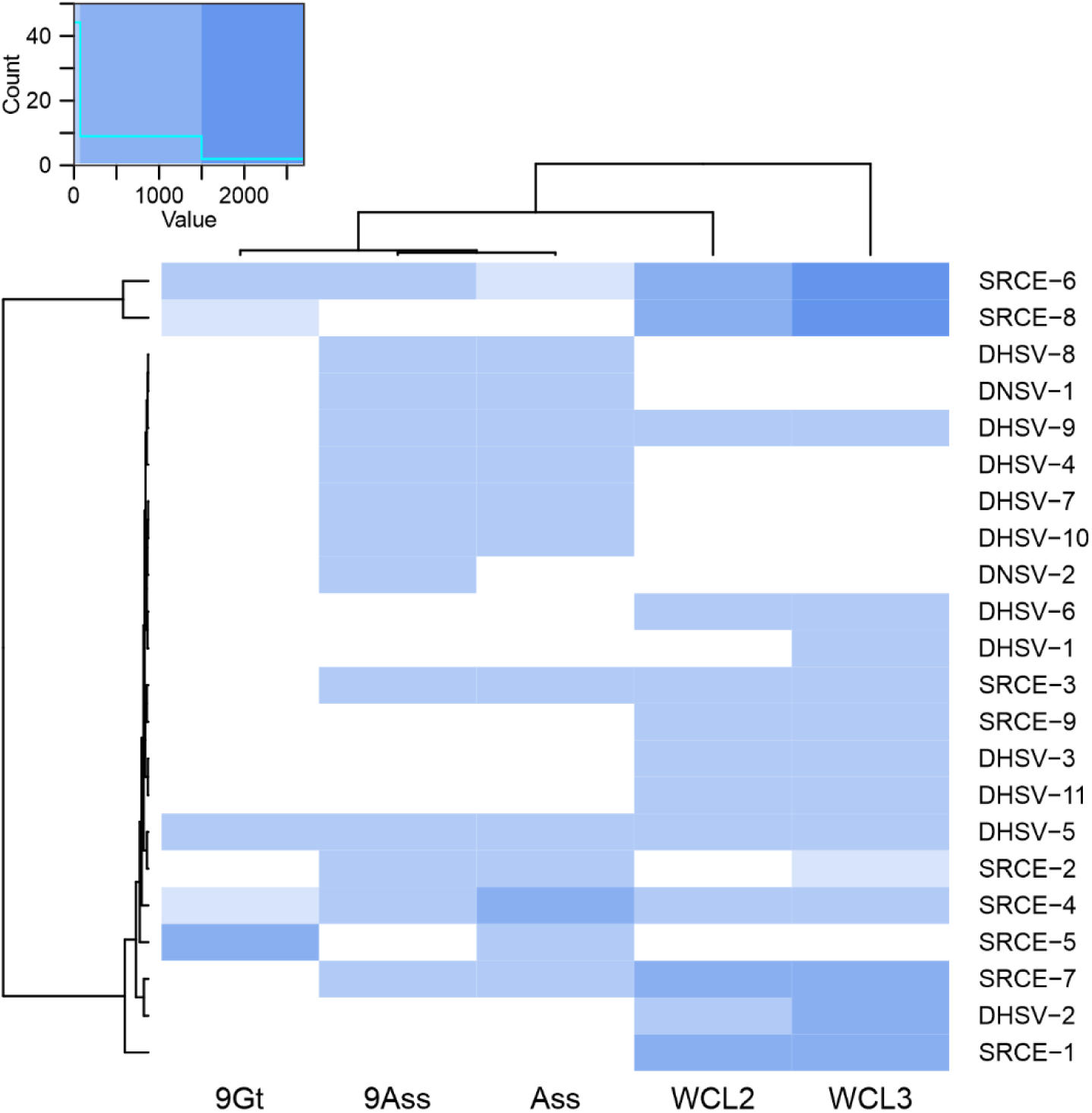
Heatmap showing the distribution and abundance of haloarchaeal and nanohaloarchaeal spindle-shaped viruses and SRCEs in Danakil salt lakes.

**Figure S12.**
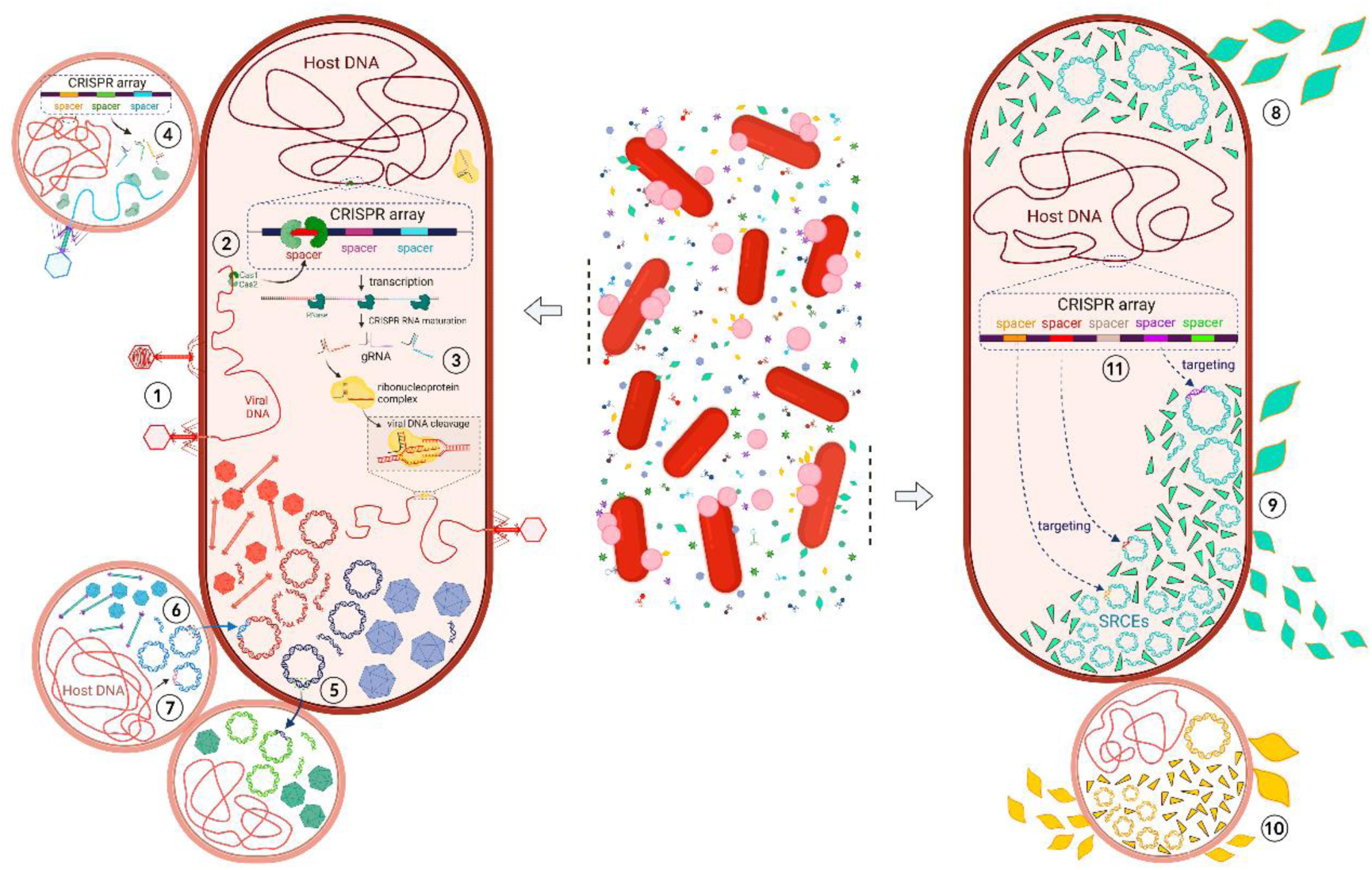
The complexity of interactions between haloarchaea, nanohaloarchaea, their respective viruses and virus satellites. 1: Viruses infect a haloarchaeal host; 2: A spacer is acquired from viral DNA by the haloarchaeal CRISPR-Cas system; 3: CRISPR spacers are transcribed, matured, and matched with target viral DNA with the help of Cas proteins, leading to the cleavage of the invading viral DNA; 4: Similar CIRSPR-Cas immunity processes (1, 2, 3) are happening in nanohaloarchaea; 5: A gene transfer from an icosahedral HV to an icosahedral NHV; 6: A gene transfer from a head-tailed NHV to a head-tailed HV; 7: A gene transfer from a nanohaloarchaeal host to a head-tailed virus; 8: Spindle-shaped HVs infect a haloarchaeal host without interference of virus satellites (SRCEs); 9: Spindle-shaped HVs and SRCEs co-infect a haloarchaeal host. SRCEs replicate and consume the virion components of HVs, which lead to a decrease in HV production; 10: Similar phenomenon (9) is also happening in nanohaloarchaea; 11: the archaeal host carries a CRISPR array with spacers targeting both viruses and virus satellites. Created in BioRender. ZHOU, Y. (2025) https://BioRender.com/z24x580

## REFERENCES

1 Jaffe, A. L. & Banfield, J. F. Candidate Phyla Radiation bacteria. Curr Biol 34, R80–R81, doi:10.1016/j.cub.2023.12.024 (2024).

2 Baker, B. J. et al. Diversity, ecology and evolution of Archaea. Nat Microbiol 5, 887–900, doi:10.1038/s41564-020-0715-z (2020).

3 Dombrowski, N., Lee, J. H., Williams, T. A., Offre, P. & Spang, A. Genomic diversity, lifestyles and evolutionary origins of DPANN archaea. FEMS Microbiol Lett 366, doi:10.1093/femsle/fnz008 (2019).

4 Rinke, C. et al. Insights into the phylogeny and coding potential of microbial dark matter. Nature 499, 431–437, doi:10.1038/nature12352 (2013).

5 Andrade, K. et al. Metagenomic and lipid analyses reveal a diel cycle in a hypersaline microbial ecosystem. ISME J 9, 2697–2711, doi:10.1038/ismej.2015.66 (2015).

6 Baker, B. A. et al. Expanded phylogeny of extremely halophilic archaea shows multiple independent adaptations to hypersaline environments. Nat Microbiol 9, 964–975, doi:10.1038/s41564-024-01647-4 (2024).

7 Feng, Y. et al. The Evolutionary Origins of Extreme Halophilic Archaeal Lineages. Genome Biol Evol 13, doi:10.1093/gbe/evab166 (2021).

8 Zhao, D. et al. Comparative Genomic Insights into the Evolution of Halobacteria-Associated “Candidatus Nanohaloarchaeota”. mSystems 7, e0066922, doi:10.1128/msystems.00669-22 (2022).

9 Narasingarao, P. et al. De novo metagenomic assembly reveals abundant novel major lineage of Archaea in hypersaline microbial communities. ISME J 6, 81–93, doi:10.1038/ismej.2011.78 (2012).

10 La Cono, V. et al. Nanohaloarchaea as beneficiaries of xylan degradation by haloarchaea. Microb Biotechnol 16, 1803–1822, doi:10.1111/1751-7915.14272 (2023).

11 La Cono, V. et al. Symbiosis between nanohaloarchaeon and haloarchaeon is based on utilization of different polysaccharides. Proc Natl Acad Sci U S A 117, 20223–20234, doi:10.1073/pnas.2007232117 (2020).

12 Hamm, J. N. et al. Unexpected host dependency of Antarctic Nanohaloarchaeota. Proc Natl Acad Sci U S A 116, 14661–14670, doi:10.1073/pnas.1905179116 (2019).

13 Reva, O. et al. Functional diversity of nanohaloarchaea within xylan-degrading consortia. Front Microbiol 14, 1182464, doi:10.3389/fmicb.2023.1182464 (2023).

14 Belilla, J. et al. Archaeal overdominance close to life-limiting conditions in geothermally influenced hypersaline lakes at the Danakil Depression, Ethiopia. Environ Microbiol 23, 7168–7182, doi:10.1111/1462-2920.15771 (2021).

15 Puxty, R. J. & Millard, A. D. Functional ecology of bacteriophages in the environment. Curr Opin Microbiol 71, 102245, doi:10.1016/j.mib.2022.102245 (2023).

16 Breitbart, M., Bonnain, C., Malki, K. & Sawaya, N. A. Phage puppet masters of the marine microbial realm. Nat Microbiol 3, 754–766, doi:10.1038/s41564-018-0166-y (2018).

17 López-García, P. et al. Metagenome-derived virus-microbe ratios across ecosystems. ISME J 17, 1552–1563, doi:10.1038/s41396-023-01431-y (2023).

18 Santos, F. et al. Culture-independent approaches for studying viruses from hypersaline environments. Appl Environ Microbiol 78, 1635–1643, doi:10.1128/AEM.07175-11 (2012).

19 Atanasova, N. S., Bamford, D. H. & Oksanen, H. M. Haloarchaeal virus morphotypes. Biochimie 118, 333–343, doi:10.1016/j.biochi.2015.07.002 (2015).

20 Luk, A. W., Williams, T. J., Erdmann, S., Papke, R. T. & Cavicchioli, R. Viruses of haloarchaea. Life (Basel) 4, 681–715, doi:10.3390/life4040681 (2014).

21 Dyall-Smith, M., Tang, S. L. & Bath, C. Haloarchaeal viruses: how diverse are they? Res Microbiol 154, 309–313, doi:10.1016/S0923-2508(03)00076-7 (2003).

22 Garcia-Heredia, I. et al. Reconstructing viral genomes from the environment using fosmid clones: the case of haloviruses. PLoS One 7, e33802, doi:10.1371/journal.pone.0033802 (2012).

23 Alarcón-Schumacher, T., Lucking, D. & Erdmann, S. Revisiting evolutionary trajectories and the organization of the *Pleolipoviridae* family. PLoS Genet 19, e1010998, doi:10.1371/journal.pgen.1010998 (2023).

24 Emerson, J. B. et al. Virus-host and CRISPR dynamics in Archaea-dominated hypersaline Lake Tyrrell, Victoria, Australia. Archaea 2013, 370871, doi:10.1155/2013/370871 (2013).

25 Emerson, J. B. et al. Dynamic viral populations in hypersaline systems as revealed by metagenomic assembly. Appl Environ Microbiol 78, 6309–6320, doi:10.1128/AEM.01212-12 (2012).

26 Crits-Christoph, A. et al. Functional interactions of archaea, bacteria and viruses in a hypersaline endolithic community. Environ Microbiol 18, 2064–2077, doi:10.1111/1462-2920.13259 (2016).

27 Martínez-García, M., Santos, F., Moreno-Paz, M., Parro, V. & Anton, J. Unveiling viral-host interactions within the ‘microbial dark matter’. Nat Commun 5, 4542, doi:10.1038/ncomms5542 (2014).

28 Wu, Z., Liu, S. & Ni, J. Metagenomic characterization of viruses and mobile genetic elements associated with the DPANN archaeal superphylum. Nat Microbiol 9, 3362–3375, doi:10.1038/s41564-024-01839-y (2024).

29 Gutiérrez-Preciado, A. et al. Extremely acidic proteomes and metabolic flexibility in bacteria and highly diversified archaea thriving in geothermal chaotropic brines. Nat Ecol Evol 8, 1856–1869, doi:10.1038/s41559-024-02505-6 (2024).

30 Atanasova, N. S., Oksanen, H. M. & Bamford, D. H. Haloviruses of archaea, bacteria, and eukaryotes. Curr Opin Microbiol 25, 40–48, doi:10.1016/j.mib.2015.04.001 (2015).

31 Sime-Ngando, T. et al. Diversity of virus-host systems in hypersaline Lake Retba, Senegal. Environ Microbiol 13, 1956–1972, doi:10.1111/j.1462-2920.2010.02323.x (2011).

32 Camargo, A. P. et al. Identification of mobile genetic elements with geNomad. Nat Biotechnol 42, 1303–1312, doi:10.1038/s41587-023-01953-y (2024).

33 Guo, J. et al. VirSorter2: a multi-classifier, expert-guided approach to detect diverse DNA and RNA viruses. Microbiome 9, 37, doi:10.1186/s40168-020-00990-y (2021).

34 Maier, L. K. et al. The nuts and bolts of the Haloferax CRISPR-Cas system I-B. RNA Biol 16, 469–480, doi:10.1080/15476286.2018.1460994 (2019).

35 Nayfach, S. et al. A genomic catalog of Earth’s microbiomes. Nat Biotechnol 39, 499–509, doi:10.1038/s41587-020-0718-6 (2021).

36 Liu, Y. et al. Diversity, taxonomy, and evolution of archaeal viruses of the class Caudoviricetes. PLoS Biol 19, e3001442, doi:10.1371/journal.pbio.3001442 (2021).

37 Baquero, D. P., Bignon, E. A. & Krupovic, M. Pleomorphic viruses establish stable relationship with marine hyperthermophilic archaea. ISME J 18, doi:10.1093/ismejo/wrae008 (2024).

38 Medvedeva, S., Borrel, G., Krupovic, M. & Gribaldo, S. A compendium of viruses from methanogenic archaea reveals their diversity and adaptations to the gut environment. Nat Microbiol 8, 2170–2182, doi:10.1038/s41564-023-01485-w (2023).

39 Han, Z. et al. Structural insights into a spindle-shaped archaeal virus with a sevenfold symmetrical tail. Proc Natl Acad Sci U S A 119, e2119439119, doi:10.1073/pnas.2119439119 (2022).

40 Wang, F. et al. Spindle-shaped archaeal viruses evolved from rod-shaped ancestors to package a larger genome. Cell 185, 1297–1307 e1211, doi:10.1016/j.cell.2022.02.019 (2022).

41 Medvedeva, S. et al. Three families of Asgard archaeal viruses identified in metagenome-assembled genomes. Nat Microbiol 7, 962–973, doi:10.1038/s41564-022-01144-6 (2022).

42 Krupovic, M., Cvirkaite-Krupovic, V., Iranzo, J., Prangishvili, D. & Koonin, E. V. Viruses of archaea: Structural, functional, environmental and evolutionary genomics. Virus Res 244, 181–193, doi:10.1016/j.virusres.2017.11.025 (2018).

43 Reed, C. J., Lewis, H., Trejo, E., Winston, V. & Evilia, C. Protein adaptations in archaeal extremophiles. Archaea 2013, 373275, doi:10.1155/2013/373275 (2013).

44 Krupovic, M., Gribaldo, S., Bamford, D. H. & Forterre, P. The evolutionary history of archaeal MCM helicases: a case study of vertical evolution combined with hitchhiking of mobile genetic elements. Mol Biol Evol 27, 2716–2732, doi:10.1093/molbev/msq161 (2010).

45 Das, G. & Varshney, U. Peptidyl-tRNA hydrolase and its critical role in protein biosynthesis. Microbiology (Reading) 152, 2191–2195, doi:10.1099/mic.0.29024-0 (2006).

46 Ho, C. K. & Shuman, S. Bacteriophage T4 RNA ligase 2 (gp24.1) exemplifies a family of RNA ligases found in all phylogenetic domains. Proc Natl Acad Sci U S A 99, 12709–12714, doi:10.1073/pnas.192184699 (2002).

47 Oakley, A. J., Coggan, M. & Board, P. G. Identification and characterization of gamma-glutamylamine cyclotransferase, an enzyme responsible for gamma-glutamyl-epsilon-lysine catabolism. J Biol Chem 285, 9642–9648, doi:10.1074/jbc.M109.082099 (2010).

48 Milewski, S. Glucosamine-6-phosphate synthase--the multi-facets enzyme. Biochim Biophys Acta 1597, 173–192, doi:10.1016/s0167-4838(02)00318-7 (2002).

49 Isupov, M. N. et al. Substrate binding is required for assembly of the active conformation of the catalytic site in Ntn amidotransferases: evidence from the 1.8 A crystal structure of the glutaminase domain of glucosamine 6-phosphate synthase. Structure 4, 801–810, doi:10.1016/s0969-2126(96)00087-1 (1996).

50 Baquero, D. P. et al. Structure and assembly of archaeal viruses. Adv Virus Res 108, 127–164, doi:10.1016/bs.aivir.2020.09.004 (2020).

51 Dedeo, C. L., Cingolani, G. & Teschke, C. M. Portal Protein: The Orchestrator of Capsid Assembly for the dsDNA Tailed Bacteriophages and Herpesviruses. Annu Rev Virol 6, 141–160, doi:10.1146/annurev-virology-092818-015819 (2019).

52 Reva, O. N. et al. DPANN symbiont of Haloferax volcanii accelerates xylan degradation by the non-host haloarchaeon Halorhabdus sp. iScience 28, 111749 (2025).

53 Arnold, H. P. et al. The genetic element pSSVx of the extremely thermophilic crenarchaeon Sulfolobus is a hybrid between a plasmid and a virus. Mol Microbiol 34, 217–226, doi:10.1046/j.1365-2958.1999.01573.x (1999).

54 Roux, S. et al. Analysis of metagenomic data reveals common features of halophilic viral communities across continents. Environ Microbiol 18, 889–903, doi:10.1111/1462-2920.13084 (2016).

55 Guyot, V. et al. A newly emerging alphasatellite affects banana bunchy top virus replication, transcription, siRNA production and transmission by aphids. PLoS Pathog 18, e1010448, doi:10.1371/journal.ppat.1010448 (2022).

56 Nasko, D. J. et al. CRISPR Spacers Indicate Preferential Matching of Specific Virioplankton Genes. mBio 10, doi:10.1128/mBio.02651-18 (2019).

57 Gao, L. A. et al. Prokaryotic innate immunity through pattern recognition of conserved viral proteins. Science 377, eabm4096, doi:10.1126/science.abm4096 (2022).

58 Krupovic, M., Dolja, V. V. & Koonin, E. V. The LUCA and its complex virome. Nat Rev Microbiol 18, 661–670, doi:10.1038/s41579-020-0408-x (2020).

59 Krupovic, M., Quemin, E. R., Bamford, D. H., Forterre, P. & Prangishvili, D. Unification of the globally distributed spindle-shaped viruses of the Archaea. J Virol 88, 2354–2358, doi:10.1128/JVI.02941-13 (2014).

60 Liu, Y., Dyall-Smith, M. & Oksanen, H. M. ICTV Virus Taxonomy Profile: Pleolipoviridae 2022. J Gen Virol 103, doi:10.1099/jgv.0.001793 (2022).

61 Esser, S. P. et al. A predicted CRISPR-mediated symbiosis between uncultivated archaea. Nat Microbiol 8, 1619–1633, doi:10.1038/s41564-023-01439-2 (2023).

62 Hadidi, A., Czosnek, H. H., Kalantidis, K. & Palukaitis, P. Viroids and Satellites and Their Vector Interactions. Viruses 16, doi:10.3390/v16101598 (2024).

63 Palukaitis, P. Satellite RNAs and Satellite Viruses. Mol Plant Microbe Interact 29, 181–186, doi:10.1094/MPMI-10-15-0232-FI (2016).

64 Francki, R. I. Plant virus satellites. Annu Rev Microbiol 39, 151–174, doi:10.1146/annurev.mi.39.100185.001055 (1985).

65 Lindqvist, B. H., Deho, G. & Calendar, R. Mechanisms of genome propagation and helper exploitation by satellite phage P4. Microbiol Rev 57, 683–702, doi:10.1128/mr.57.3.683-702.1993 (1993).

66 Boyd, C. M. & Seed, K. D. A phage satellite manipulates the viral DNA packaging motor to inhibit phage and promote satellite spread. Nucleic Acids Res 52, 10431–10446, doi:10.1093/nar/gkae675 (2024).

67 Barcia-Cruz, R. et al. Phage-inducible chromosomal minimalist islands (PICMIs), a novel family of small marine satellites of virulent phages. Nat Commun 15, 664, doi:10.1038/s41467-024-44965-1 (2024).

68 de Sousa, J. A. M., Fillol-Salom, A., Penades, J. R. & Rocha, E. P. C. Identification and characterization of thousands of bacteriophage satellites across bacteria. Nucleic Acids Res 51, 2759–2777, doi:10.1093/nar/gkad123 (2023).

69 Hackl, T. et al. Novel integrative elements and genomic plasticity in ocean ecosystems. Cell 186, 47–62 e16, doi:10.1016/j.cell.2022.12.006 (2023).

70 Alqurainy, N. et al. A widespread family of phage-inducible chromosomal islands only steals bacteriophage tails to spread in nature. Cell Host Microbe 31, 69–82 e65, doi:10.1016/j.chom.2022.12.001 (2023).

71 Ibarra-Chavez, R., Hansen, M. F., Pinilla-Redondo, R., Seed, K. D. & Trivedi, U. Phage satellites and their emerging applications in biotechnology. FEMS Microbiol Rev 45, doi:10.1093/femsre/fuab031 (2021).

72 Wang, Y. et al. A novel Sulfolobus non-conjugative extrachromosomal genetic element capable of integration into the host genome and spreading in the presence of a fusellovirus. Virology 363, 124–133, doi:10.1016/j.virol.2007.01.035 (2007).

73 Dyall-Smith, M. & Pfeiffer, F. Global Distribution and Diversity of Haloarchaeal pL6-Family Plasmids. Genes (Basel) 15, doi:10.3390/genes15091123 (2024).

74 Dyall-Smith, M. & Pfeiffer, F. The PL6-Family Plasmids of Haloquadratum Are Virus-Related. Front Microbiol 9, 1070, doi:10.3389/fmicb.2018.01070 (2018).

75 Allers, T., Ngo, H. P., Mevarech, M. & Lloyd, R. G. Development of additional selectable markers for the halophilic archaeon Haloferax volcanii based on the leuB and trpA genes. Appl Environ Microbiol 70, 943–953, doi:10.1128/AEM.70.2.943-953.2004 (2004).

76 Zhou, Y., Wang, Y., Prangishvili, D. & Krupovic, M. Exploring the Archaeal Virosphere by Metagenomics. Methods Mol Biol 2732, 1–22, doi:10.1007/978-1-0716-3515-5_1 (2024).

77 Nayfach, S. et al. CheckV assesses the quality and completeness of metagenome-assembled viral genomes. Nat Biotechnol 39, 578–585, doi:10.1038/s41587-020-00774-7 (2021).

78 Bin Jang, H., et al. Taxonomic assignment of uncultivated prokaryotic virus genomes is enabled by gene-sharing networks. Nat Biotechnol 37, 632–639, doi:10.1038/s41587-019-0100-8 (2019).

79 Shannon, P. et al. Cytoscape: a software environment for integrated models of biomolecular interaction networks. Genome Res 13, 2498–2504, doi:10.1101/gr.1239303 (2003).

80 Couvin, D. et al. CRISPRCasFinder, an update of CRISRFinder, includes a portable version, enhanced performance and integrates search for Cas proteins. Nucleic Acids Res 46, W246–W251, doi:10.1093/nar/gky425 (2018).

81 Li, W. & Godzik, A. Cd-hit: a fast program for clustering and comparing large sets of protein or nucleotide sequences. Bioinformatics 22, 1658–1659, doi:10.1093/bioinformatics/btl158 (2006).

82 Zhou, Y. et al. Diverse viruses of marine archaea discovered using metagenomics. Environ Microbiol 25, 367–382, doi:10.1111/1462-2920.16287 (2023).

83 Edwards, R. A., McNair, K., Faust, K., Raes, J. & Dutilh, B. E. Computational approaches to predict bacteriophage-host relationships. FEMS Microbiol Rev 40, 258–272, doi:10.1093/femsre/fuv048 (2016).

84 Gong, C. et al. Novel Virophages Discovered in a Freshwater Lake in China. Front Microbiol 7, 5, doi:10.3389/fmicb.2016.00005 (2016).

85 Seemann, T. Prokka: rapid prokaryotic genome annotation. Bioinformatics 30, 2068–2069, doi:10.1093/bioinformatics/btu153 (2014).

86 Steinegger, M. et al. HH-suite3 for fast remote homology detection and deep protein annotation. BMC Bioinformatics 20, 473, doi:10.1186/s12859-019-3019-7 (2019).

87 Gabler, F. et al. Protein Sequence Analysis Using the MPI Bioinformatics Toolkit. Curr Protoc Bioinformatics 72, e108, doi:10.1002/cpbi.108 (2020).

88 Nishimura, Y. et al. ViPTree: the viral proteomic tree server. Bioinformatics 33, 2379–2380, doi:10.1093/bioinformatics/btx157 (2017).

89 Aroney, S. T., et al. CoverM: Read alignment statistics for metagenomics. *arXiv* 10.48550/arXiv.2501.11217 (2025).

90 Edgar, R. C. Muscle5: High-accuracy alignment ensembles enable unbiased assessments of sequence homology and phylogeny. Nat Commun 13, 6968, doi:10.1038/s41467-022-34630-w (2022).

91 Capella-Gutiérrez, S., Silla-Martínez, J. M. & Gabaldon, T. trimAl: a tool for automated alignment trimming in large-scale phylogenetic analyses. Bioinformatics 25, 1972–1973, doi:10.1093/bioinformatics/btp348 (2009).

92 Minh, B. Q. et al. IQ-TREE 2: New Models and Efficient Methods for Phylogenetic Inference in the Genomic Era. Mol Biol Evol 37, 1530–1534, doi:10.1093/molbev/msaa015 (2020).

93 Letunic, I. & Bork, P. Interactive Tree of Life (iTOL) v6: recent updates to the phylogenetic tree display and annotation tool. Nucleic Acids Res 52, W78–W82, doi:10.1093/nar/gkae268 (2024).

94 Kazlauskas, D., Varsani, A., Koonin, E. V. & Krupovic, M. Multiple origins of prokaryotic and eukaryotic single-stranded DNA viruses from bacterial and archaeal plasmids. Nat Commun 10, 3425, doi:10.1038/s41467-019-11433-0 (2019).

## SUPPLEMENTARY REFERENCES

1 Gross, M., Marianovsky, I. & Glaser, G. MazG -- a regulator of programmed cell death in Escherichia coli. Mol Microbiol 59, 590–601, doi:10.1111/j.1365-2958.2005.04956.x (2006).

2 Liu, Y. et al. Diversity, taxonomy, and evolution of archaeal viruses of the class Caudoviricetes. PLoS Biol 19, e3001442, doi:10.1371/journal.pbio.3001442 (2021).

3 Philosof, A. et al. Novel Abundant Oceanic Viruses of Uncultured Marine Group II Euryarchaeota. Curr Biol 27, 1362–1368, doi:10.1016/j.cub.2017.03.052 (2017).

4 Zhou, Y. et al. Diverse viruses of marine archaea discovered using metagenomics. Environ Microbiol 25, 367–382, doi:10.1111/1462-2920.16287 (2023).

5 Gutiérrez-Preciado, A. et al. Extremely acidic proteomes and metabolic flexibility in bacteria and highly diversified archaea thriving in geothermal chaotropic brines. Nat Ecol Evol 8, 1856–1869, doi:10.1038/s41559-024-02505-6 (2024).

